# Efficient long fragment editing technique enables large-scale and scarless bacterial genome engineering

**DOI:** 10.1101/2020.01.28.922609

**Authors:** Chaoyong Huang, Liwei Guo, Jingge Wang, Ning Wang, Yi-Xin Huo

## Abstract

Bacteria are versatile living systems that enhance our understanding of nature and enable biosynthesis of valuable chemicals. Long fragment editing techniques are of great importance for accelerating bacterial genome engineering to obtain desirable and genetically stable strains. However, the existing genome editing methods cannot meet the needs of engineers. We herein report an efficient long fragment editing method for large-scale and scarless genome engineering in *Escherichia coli*. The method enabled us to insert DNA fragments up to 12 kb into the genome and to delete DNA fragments up to 186.7 kb from the genome, with positive rates over 95%. We applied this method for *E. coli* genome simplification, resulting in 12 individual deletion mutants and four cumulative deletion mutants. The simplest genome lost a total of 370.6 kb of DNA sequence containing 364 open reading frames. Additionally, we applied this technique to metabolic engineering and obtained a genetically stable plasmid-independent isobutanol production strain that produced 1.3 g/L isobutanol via shake-flask fermentation. These results suggest that the method is a powerful genome engineering tool, highlighting its potential to be applied in synthetic biology and metabolic engineering.

## 1 Introduction

As a class of versatile living systems, bacteria are useful in many fields of synthetic biology. In bacteria, genetic information contained on the single-copy genome determines the characteristics of a specific strain. To understand bacterial characteristics and utilize them to explore the world and serve human life, researchers frequently conduct genome engineering to reprogram the genetic information of bacteria. Through DNA editing, researchers can add desired exogenous genetic information to or delete unwanted endogenous genetic information from the bacterial genome. The long fragment editing technique is of great importance in accelerating bacterial genome engineering to obtain genetically stable strains. For example, the long fragment deletion technique can help to simplify the bacterial genome to explore the minimal genome of a specific strain [1, 2], and the long fragment insertion technique can help to expand the bacterial genome to archive the expanding information of the human world [3]. In metabolic engineering, plasmid maintenance requires continuous antibiotic use, which has led to biosafety issues and elevated industrial cost [4]. The long fragment editing technique is an ideal tool for constructing plasmid-independent and high-production strains.

To accelerate the process of genome engineering, researchers have developed many methods for generating insertions and deletions in the bacterial genome. Homologous recombination with polymerase chain reaction (PCR) fragments forms the basis of these methods [5, 6]. However, since RecA-mediated homologous recombination with linear DNA is of low efficiency, researchers created the desired mutagenesis on a suitable plasmid before recombining it into the genome [7-9]. To enhance the efficiency of homologous recombination, the bacteriophage-derived λ-Red system was introduced into bacteria on either the genome or plasmids. Genome editing based on λ-Red recombinases is referred to as recombineering [10-12]. In recombineering, an antibiotic resistance gene is required as a selectable marker. To remove the selectable marker after genome editing, researchers introduced counter-selection systems or site-specific recombination systems, including FLP/*FRT* and Cre/*loxP* [13, 14]. Though recombineering can handle the insertion and deletion of short DNA fragments [15-17], the editing efficiency decreases dramatically for long fragments [11]. Moreover, eliminating selectable markers and plasmids is complicated and time-consuming, and the residual *FRT* or *loxP* site may influence a later round of genome editing [13]. Generating a double-strand break (DSB) in the target DNA is an effective strategy for improving the efficiency of long fragment editing. Though the homing endonuclease I-*Sce*I is efficient for cleaving double-stranded DNA (dsDNA), researchers had to integrate an 18-bp recognition site into the target DNA before inducing DNA cleavage [18-20]. Recently, clustered regularly interspaced short palindromic repeats (CRISPR)/CRISPR-associated protein 9 (Cas9) technology was developed based on research into the adaptive immune system of *Streptococcus pneumoniae* [21]. Cas9 endonuclease complexed with a designed single-guide RNA (sgRNA) can generate DSB in a specific protospacer sequence where a proper protospacer-adjacent motif (PAM) exists [21-23]. The technique relies on sgRNA-directed cleavage at the target site to kill wild-type cells, thus circumventing the need for selectable markers or counter-selection systems. In addition, changing the 20-bp spacer sequence can reprogram the specificity of the Cas9-sgRNA complex. In the past seven years, many genome editing methods and protocols based on the CRISPR/Cas9 technique have been reported. These methods are efficient for short fragment editing in bacteria, however, for long fragment editing, significant work needs to be done.

Among all CRISPR/Cas9-based genome editing methods, CRISPR/Cas9-assisted recombineering methods perform very well. Existing CRISPR/Cas9-assisted recombineering methods use cycle DNA (plasmid-borne dsDNA) [24-26] or linear DNA (PCR-amplified dsDNA or synthesized ssDNA) [23, 24, 27-29] as the editing template, and both kinds of templates have advantages and disadvantages. The cycle editing template can avoid the attack by DNA exonucleases and copy itself along with the plasmid replication, thus resulting in much higher homologous recombination efficiency and editing efficiency. However, there is a significant chance that the total plasmid will be integrated into the genome, and these recombination events are difficult to distinguish from desired recombination events through conventional PCR verification, leading to high false positive rates. It seems that this phenomenon has not yet been noticed. A linear editing template circumvents the trouble caused by plasmid integration, thus resulting in a higher positive rate. However, the sensitivity of the linear editing template to DNA exonucleases leads to lower homologous recombination efficiency and editing efficiency. To solve this contradiction, we made a systematic optimization in the basis of existing CRISPR/Cas9-assisted recombineering methods and developed an efficient long fragment editing method for large-scale and scarless genome engineering in *Escherichia coli.* This method enabled us to insert and delete large DNA fragments into and from the genome, with high positive rates and editing efficiency. Notably, the high performance of the method was independent of high transformation efficiency, making the method much easier to put into practice. Furthermore, the method was successfully applied in genome simplification and metabolic engineering, demonstrating its value as a genome engineering tool for constructing genetically stable *E. coli* strains. We believe that this method has the potential to be widely used in other sequenced organisms in addition to *E. coli*.

## 2 Material and methods

### 2.1 Strains and culture conditions

*E. coli* strain DH5α (American Type Culture Collection, ATCC 68233) served as the host strain for molecular cloning and plasmid manipulation. MG1655 (ATCC 47076) served as the genetic material in editing experiments except where otherwise stated. The strains involved in this study are listed in Table S1. The verification primers used in the genome editing experiments are listed in Table S2. Luria-Bertani (LB) medium (10 g/L tryptone, 5 g/L yeast extract, and 10 g/L NaCl) was used for cell growth in all cases except where otherwise noted. The solid medium contained 20 g/L agar. Super Optimal broth with Catabolite repression (SOC) medium (20 g/L tryptone, 5 g/L yeast extract, 0.5 g/L NaCl, 2.5 mM KCl, 10 mM MgCl_2_, 10 mM MgSO_4_, and 20 mM glucose) was used for cell recovery. M9 medium (6 g/L Na_2_HPO_4_, 3 g/L KH_2_PO_4_, 0.5 g/L NaCl, 1 g/L NH_4_Cl, 1 mM MgSO_4_, 0.1 mM CaCl_2_, 10 mg/L VB_1_, 40 g/L glucose, and 4 g/L yeast extract) was used for shake-flask fermentation. The working concentrations of ampicillin (Amp) and kanamycin (Kan) were 0.1 g/L and 0.025 g/L, respectively. The working concentrations of isopropyl-β-D-thiogalactopyranoside (IPTG), X-gal, glucose, and sucrose in media or cultures were 1 mM, 0.1 g/L, 10 g/L, and 20 g/L, respectively. The working concentration of L-arabinose was 20 mM in liquid media and 5 mM in solid media. Details of the reagents and media used in this study are listed in Table S3.

### 2.2 Plasmid construction

The plasmids involved in this study are listed in Table S4. The complete sequences of plasmids p15A-P_araB_-Cas9-P_T5_-Redγβα, pSC101-P_araB_-sgRNA-Donor-T1, pSC101-P_araB_-sgRNA-Donor-T2, and pSC101-P_araB_-sgRNA-Donor-T3 are presented in Notes S1–S4. The CRISPR target sequences designed in this study are listed in Table S5. The construction of plasmid pSC101-P_araB_-sgRNA-Donor was the key step in the genome editing experiments. When constructing the pSC101-P_araB_-sgRNA-Donor plasmid containing one sgRNA expression chimera, pSC101-P_araB_-sgRNA-Donor-T1 served as the parental plasmid. First, a specifically designed donor DNA was integrated into pSC101-P_araB_-sgRNA-Donor-T1 to construct an intermediate plasmid. The donor DNA contained two homologous arms of approximately 500 bp. Then, a specific spacer (20 bp) was inserted into the intermediate plasmid between the *araB* promoter and the gRNA scaffold via single PCR and single Gibson Assembly [30]. The spacer introduced by PCR served as the overlap in Gibson Assembly. When constructing the pSC101-P_araB_-sgRNA-Donor plasmid containing two sgRNA expression chimeras, pSC101-P_araB_-sgRNA-Donor-T2 and pSC101-P_araB_-sgRNA-Donor-T3 served as the parental plasmids. First, a specifically designed donor DNA was integrated into pSC101-P_araB_-sgRNA-Donor-T2 to construct an intermediate plasmid. Then, the intermediate plasmid and pSC101-P_araB_-sgRNA-Donor-T3 were combined to construct the pSC101-P_araB_-sgRNA-Donor plasmid through PCR and Gibson Assembly. The two specific spacers introduced by PCR served as overlaps in Gibson Assembly. The detailed construction procedures of the pSC101-P_araB_-sgRNA-Donor plasmid are illustrated in Fig. S1. To reduce construction procedures, the plasmid pSC101-P_araB_-sgRNA-Donor can also be obtained through single Gibson Assembly of multiple fragments.

### 2.3 Procedures for genome editing, plasmid curing, and iterative editing

First, the Kan-resistant (Kan^R^) plasmid p15A-P_araB_-Cas9-P_T5_-Redγβα (plasmid#1) was transformed into the target strain such as MG1655 to obtain the corresponding transformants such as MG1655/plasmid#1. A series of temperature-sensitive Amp-resistant (Amp^R^) plasmids were constructed to express specific sgRNA and generate specific donor DNA, and these plasmids were collectively named pSC101-P_araB_-sgRNA-Donor (plasmid#2). Then, specific plasmid#2 was transformed into the MG1655/plasmid#1 strain, and the MG1655/plasmid#1/plasmid#2 strain was screened in a LB plate with Amp, Kan, and glucose at 30 °C. One or several single colonies were inoculated into 2 mL LB medium, and the culture was cultivated at 30 °C for 2 h (Time 1). Then, 2 μL Amp, 2 μL Kan, and 20 μL IPTG were added to the culture. After 1 h (Time 2), 20 μL L-arabinose was added, and the cultures were cultivated for another 3 h (Time 3) before plating. A 1-μL or 0.1-μL aliquot of the culture was plated onto a LB plate containing Amp, Kan, and L-arabinose, and the plate was incubated overnight at 30 °C. Positive mutants were verified by colony PCR and sequencing. The flowchart of genome editing is shown in Fig. 1 and Fig. S2. The positive mutant was cultivated in LB medium in the presence of only Kan at 40 °C for 12 h to remove the temperature-sensitive Amp^R^ plasmid#2 (Fig. S3a). Then, the obtained edited strain containing only plasmid#1 was used as the starting strain for the next round of genome editing. The Kan^R^ plasmid#1 is not stable in the host strain in the absence of Kan. When the final round of genome editing was completed, the edited strain was cultivated in LB antibiotic-free medium at 40 °C for 12 h to remove both the Amp^R^ plasmid#2 and the sucrose-sensitive Kan^R^ plasmid#1 (Fig. S3b). The overnight culture was diluted for plating on a LB plate containing sucrose. Theoretically, colonies grown on the plate are plasmid-free. For further verification, single colonies were inoculated into LB medium with or without corresponding antibiotics. The flowchart of plasmid curing and iterative editing is shown in Fig. S2.

**Fig.1.**
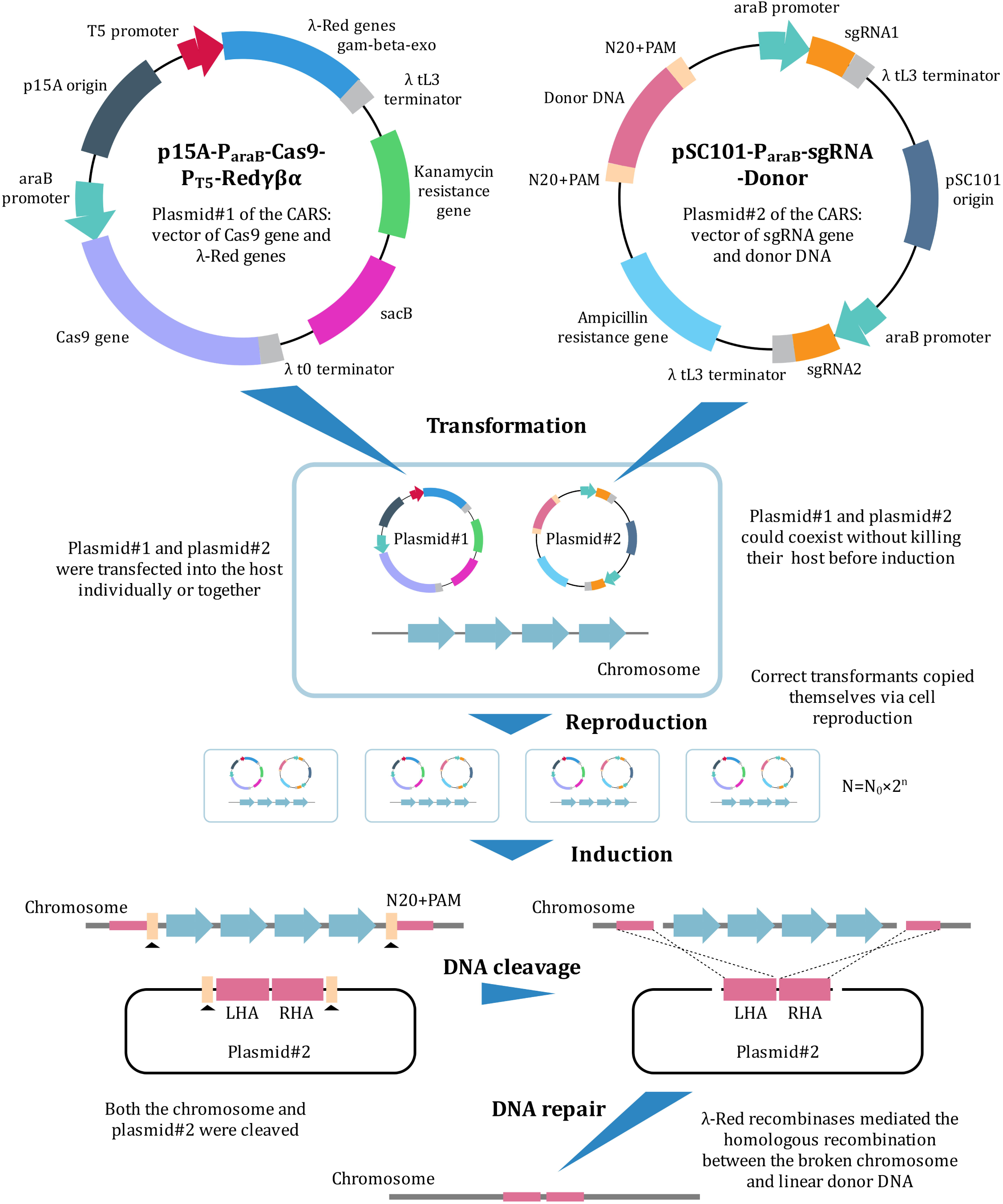
Constitutions of the genome editing system and schematic of genome editing. LHA: left homologous arm. RHA: right homologous arm.

### 2.4 Calculation of positive rate and editing efficiency

One hundred colonies in the LB plate containing Amp, Kan, and L-arabinose were tested by colony PCR to screen for positive mutants. Twenty of the positive mutants were further verified via sequencing. The positive rate was calculated as the proportion of positive colonies to the total number of colonies. In blue-white selection experiments, positive colonies were also recognized by their color. White colonies were positive, and blue colonies were negative. One control group was set along with the experimental group to calculate editing efficiency. In the control group, L-arabinose was not added, and thus no Cas9 protein and sgRNA were expressed. All other conditions and processes were the same as for the experimental group. The editing efficiency was calculated as the proportion of positive colonies in the experimental group to the total number of colonies in the control group.

### 2.5 Measurement of growth curve and transformation efficiency

For measuring the growth curve, one single colony was inoculated into 5 mL LB medium, and the culture was cultivated at 37 °C for 12 h. Then, 1 mL seed liquid was inoculated into 100 mL fresh LB medium, and the culture was cultivated at 37 °C in a 220-rpm shaker. During the 12-h cultivation, samples were taken every hour to measure the optical density of the culture at a wavelength of 600 nm (OD600) using an ultraviolet spectrophotometer (V-5100, Shanghai Metash Instruments Co., Ltd). For measuring transformation efficiency, pure pUC19 was used as supercoiled DNA. First, 1 μL pUC19 (1 ng/μL) was added to one tube of competent cells (100 μL). Next, the mixture was incubated for 30 min before conducting heat-shock for 1 min in a 42 °C water bath. Then, the tube was placed on ice for 2 min before adding 900 μL 37 °C SOC medium, and the tube was shaken at 200–230 rpm (37 °C) for 40 min. Finally, 100 μL of the cultures was plated on a LB plate containing Amp, and the plate was incubated overnight at 37 °C. The transformation efficiency is *N* × 10^4^ CFU/μg pUC19 (“*N*” refers to the number of transformants obtained in the plate).

### 2.6 Shake-flask fermentation and product detection

For testing isobutanol production, single colonies of engineered strains were inoculated into 5 mL LB media containing the appropriate antibiotics, and the cultures were cultivated at 37 °C for 12 h. Then, 200-μL seed liquid was transferred to airtight shake flasks containing 20 mL antibiotic-free M9 medium for micro-aerobic fermentation. During the 72-h fermentation, samples were taken every 12 h to test the biomass and the titer of isobutanol. Biomass was evaluated by measuring the OD600 of fermentation broth with an ultraviolet spectrophotometer (V-5100, Shanghai Metash Instruments Co., Ltd). For measuring isobutanol concentration, the fermentation broth was centrifuged at 1400 × *g* for 10 min. The supernatant was tested using a gas chromatograph (PANNA GCA91, Shanghai Wangxu Electric Co., Ltd) with high-purity isobutanol as the standard and high-purity n-pentanol as an internal reference.

## 3 Results

### 3.1 Optimization of CRISPR/Cas9-assisted recombineering method

We constructed a two-plasmid genome editing system that consists of five elements: A Cas9-expressing cassette; an sgRNA-expressing cassette; a λ-Red recombination system; a donor DNA-generation system; and a plasmid curing system (Fig. 1). A two-plasmid system is more convenient in plasmid construction than a one-plasmid system [25, 27]. For the convenience of description, the two plasmids are referred to as plasmid#1 and plasmid#2, respectively, in the following text. Specifically, Cas9 protein and λ-Red recombinases (Gam, Beta, and Exo) were expressed by plasmid#1, which contained a p15A replication origin and a Kan^R^ gene. Targeting sgRNA was expressed by plasmid#2, which contained a pSC101 replication origin and an Amp^R^ gene (Fig. 1). There were two types of plasmid#2, the first containing two sgRNA-expressing cassettes, and the second containing one sgRNA-expression cassette. The variant of plasmid#2 depended on the type of genome editing. Donor DNA, which served as an editing template to introduce sequence deletions, insertions, or replacements, was integrated into plasmid#2 to circumvent exonucleases’ attack and copy itself along with plasmid replication. The target site (N20 + PAM) on the genome was added to plasmid#2 in the flanks of donor DNA, thus the Donor DNA was released from the plasmid#2 by Cas9 cleavage during genome editing. The generated linear editing template participated in homologous recombination with the cleaved genome DNA (Fig. 1).

An inducible promoter was used to control the expressions of both Cas9 and sgRNA, thus the CRISPR/Cas9 system functioned only when an inducer was added. The L-arabinose-induced P_araB_ promoter and two IPTG-induced promoters, namely P_T5_ and P_L_lacO_1_, were tested individually (Table S6). As the P_araB_ promoter was stricter than both P_T5_ and P_L_lacO_1_, it was used for the expressions of Cas9 and sgRNA. Similarly, an inducible promoter was used to control the expressions of λ-Red recombinases, thus the λ-Red recombination system functioned only when needed, as continuous expression of λ-Red recombinases may lead to genome instability. The promoters P_araB_, P_T5_, and P_L_lacO_1_ were tested individually, and the P_T5_ promoter performed best (Table S6). In addition, we also optimized other terms, including the concentrations of L-arabinose and IPTG, the culture time, and the medium to further improve the system’s performance (Table S6). At an appropriate concentration of L-arabinose, the expression levels of Cas9 and sgRNA were enough for cleaving the single-copy genome, but insufficient for cleaving all copies of plasmid#2 (about five copies [31]). Therefore, edited cells still possessed resistance to Amp. To construct the plasmid curing system, we used the temperature-sensitive pSC101 replication origin for plasmid#2 and added the sucrose-sensitive *sacB* gene to plasmid#1 as a counter-selection marker (Fig. 1).

Each cycle of editing started with the transfection of plasmid#2 into cells containing plasmid#1 (Fig. 1 and Fig. S2). Then, the correct transformants containing the two plasmids were cultivated for cell reproduction before adding inducers to trigger DNA cleavage and DSB repair. Theoretically, sgRNA guides Cas9 to recognize and cleave the target DNA, generating DSB in the genome and plasmid#2. Then, the λ-Red recombinases mediate homologous recombination between the broken genome and linear donor DNA. This transfers the desired mutation from the donor DNA to the genome, destroying the target site (Fig. 1 and Fig. S2). The cells acquiring the desired mutation survive, and these with an unrepaired genome undergo cell death. Thus, plating liquid cultures on agar medium containing Kan and Amp allowed the selection of desired clones. Colonies growing on the plates were further verified through PCR and sequencing. Then, correct mutants were cultivated at 40 °C in medium containing only Kan to eliminate plasmid#2 (Fig. S3a). The cultures were inoculated into fresh medium to prepare competent cells for a new round of editing (Fig. S2). Each cycle of editing required three days. After the final round of editing, plasmid#1 and plasmid#2 were eliminated together by incubating the correct clones at 40 °C in antibiotic-free medium and plating the cultures on agar medium containing sucrose (Fig. S2 and Fig. S3).

### 3.2 Long fragment insertion

To evaluate the ability of the genome editing method to mediate long fragment insertion, we tried to insert fragments of different lengths (3 kb, 6 kb, 9 kb, and 12 kb) into the *lacZ* gene of *E. coli* strain MG1655 (Fig. 2a). We constructed four different versions of plasmid#2 harboring the corresponding donor DNA and expressing the same sgRNA targeting the *lacZ* gene. The four inserted fragments came from the F plasmid of *E. coli* strain XL1-Blue, and they had no homology with the MG1655 genome. The insertion of these fragments would inactivate the *lacZ* gene encoding β-galactosidase. Thus, we could differentiate edited and unedited colonies via blue-white selection. The edited colonies were white in a LB plate containing IPTG and X-gal, while the unedited colonies were blue. We also identified edited clones though PCR. One pair of primers (F1/R1) was designed for the verification of 3-kb insertion (Fig. 2a), and correct clones obtained much larger PCR products than the control (Fig. S4a). Two pairs of primers were designed for the verification of 6-kb, 9-kb, and 12-kb insertions (Fig. 2a). The correct clones obtained the desired PCR products using both F1/R2 and F2-X/R1 (X=1, 2, 3), while the control did not (Fig. S4b–d). The PCR products were further verified by sequencing. Based on the results of blue-white selection, PCR, and sequencing, we determined the editing efficiencies and positive rates. The editing efficiencies in these four insertion experiments were 1.2 × 10^−3^, 1.2 × 10^−3^, 9.6 × 10^−4^, and 7.2 × 10^−4^, respectively (Fig. 2b). The positive rates in the four insertion experiments were 97.3%, 98.3%, 96.7%, and 98.3%, respectively (Fig. 2b). These results indicated that both Cas9-mediated DNA cleavage and λ-Red-mediated DSB repair were efficient in our experiments. We found that the small-proportion negative colonies (< 5%), commonly called “escapers” [23, 27], came from two sources. More than half of the escapers did not undergo cleavage by Cas9, probably because of the limited induction time and intensity of L-arabinose. The remaining escapers acquired deletions of unknown length in the target site, which was likely due to the presence of A-EJ repair [32, 33]. The 12-kb insertion is sufficient for application in most cases of synthetic biology and metabolic engineering. The method was compared to three existing CRISPR/Cas9-based methods that performed relatively well in long fragment insertion (Fig. 2c). Method 1 enabled the insertion of 8-kb exogenous DNA, yielding a positive rate of 15% [27]. Method 2 enabled the insertion of 7-kb exogenous DNA, and the positive rate was 61% in the presence of a selectable marker [34]. Method 3 enabled the insertion of 7-kb exogenous DNA, yielding a positive rate of 10% [35]. Our method performed better as it enabled the insertion of 12-kb exogenous DNA with a positive rate of 98.3%.

**Fig.2.**
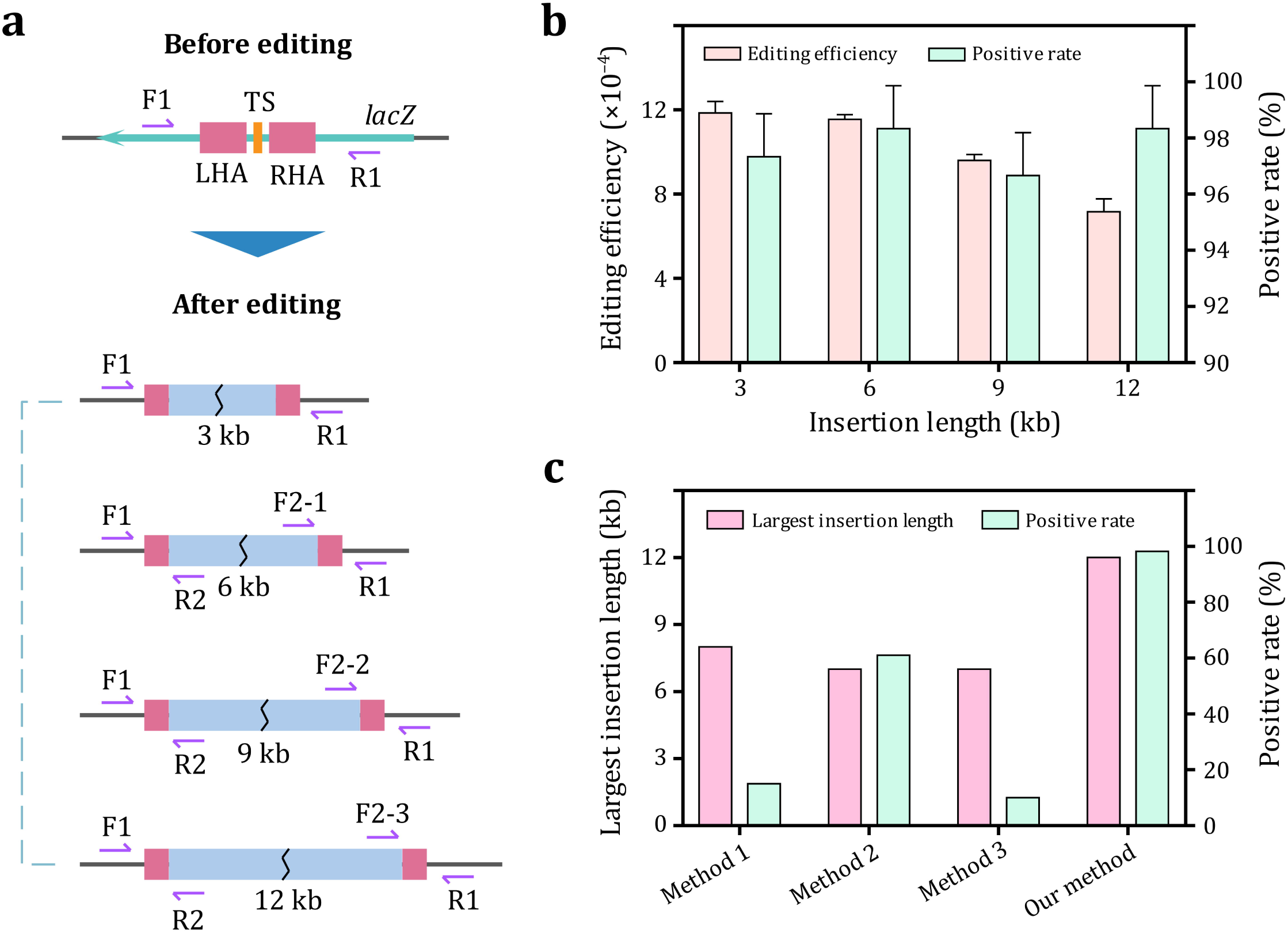
Long fragment insertion. (**a**) Schematic of fragment insertion of different lengths. TS: target site. LHA: left homologous arm. RHA: right homologous arm. F: forward primer. R: reverse primer. (**b**) Editing efficiencies and positive rates in the four editing experiments. (**c**) Comparison of largest insertion length and positive rate between three reported methods and our method. Data are expressed as means ± s.d. from three independent experiments.

### 3.3 Long fragment deletion

First, we successfully deleted a 99.9-kb fragment, starting at 565,156 and ending at 665,088, in the MG1655 genome (Fig. 3a). To determine the relationship between editing performance and the length of the deleted fragment, we selected seven fragments of different lengths within the 99.9-kb fragment for individual deletion. The lengths of these fragments were 9.1 kb, 21.5 kb, 30.6 kb, 39.4 kb, 59.8 kb, 79.8 kb, and 99.9 kb (Fig. 3a). To delete these fragments, we constructed seven different versions of plasmid#2 harboring two sgRNA-expressing cassettes. One sgRNA targets the same site (TS1) in the genome, and the other targets different sites (TS2-1–TS2-7) (Fig. 3a). Based on the results of PCR and sequencing, we determined their editing efficiencies and positive rates (Fig. 3b). As demonstrated, all positive rates were over 95%, similar to the results in long fragment insertion experiments. The deletion of 9.1-kb, 21.5-kb, 30.6-kb, 39.4-kb, 59.8-kb, and 79.8-kb fragments resulted in similar editing efficiencies, while the deletion of the 99.9-kb fragment resulted in lower editing efficiency (Fig. 3b). We found that the 99.9-kb fragment knockout strain grew much more slowly than wild-type MG1655, while the 79.8-kb fragment knockout strain had a similar growth rate to wild-type MG1655 (Fig. S5a and S5d). This phenomenon implies that the terminal region of the 99.9-kb fragment contained some genetic information that was important for cell growth. The decrease in editing efficiency of the 99.9-kb deletion experiment might be due to the lower viability of edited cells, as the editing efficiency was calculated based on the number of colonies formed on the plate. In this study, we also successfully deleted other long fragments in the genome (Fig. 4d), which are described in the next section. The method was compared to four existing CRISPR/Cas9-based methods that performed relatively well in long fragment deletion (Fig. 3c). Method 1 enabled the deletion of 12 kb of genome DNA, yielding a positive rate of 90% [27]. Method 2 enabled the deletion of 17 kb of genome DNA, and the positive rate was 17% [36]. Method 3 enabled the deletion of 100 kb of genome DNA, yielding a positive rate of 75% [28]. Method 4 enabled the deletion of 123 kb of genome DNA, and the positive rate was 36% [37]. Our method performed better as it enabled the deletion 186.7 kb of genome DNA with a positive rate of 96.6%.

**Fig.3.**
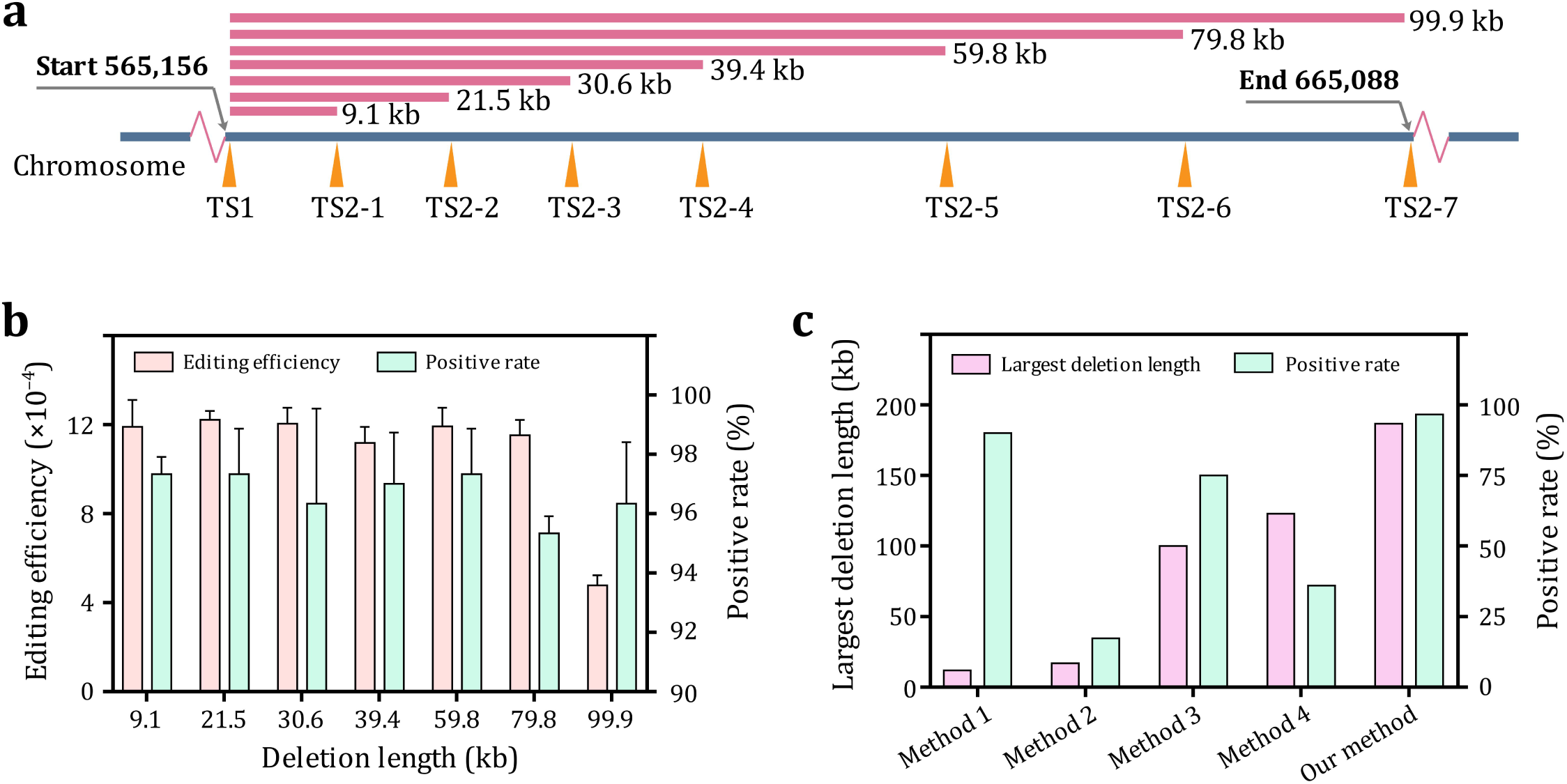
Long fragment deletion. (**a**) Schematic of fragment deletion of different lengths. TS: target site. (**b**) Editing efficiencies and positive rates in the seven editing experiments. (**c**) Comparison of largest deletion length and positive rate between four reported methods and our method. Data are expressed as means ± s.d. from three independent experiments.

**Fig.4.**
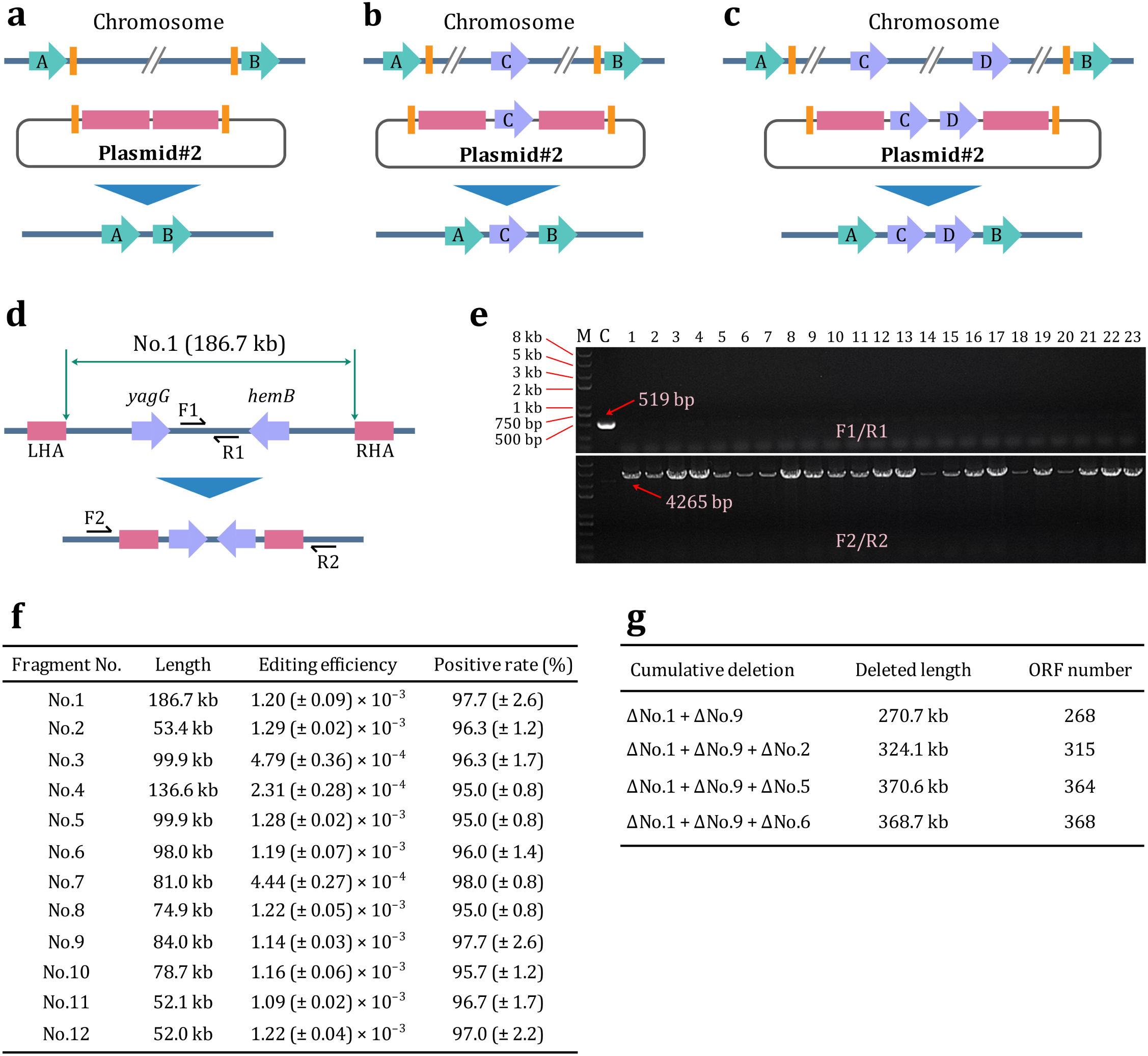
Deletion of nonessential sequences and genome simplification. (**a**) Deletion of long fragments containing no essential gene. (**b**) Deletion of long fragments containing one essential gene. (**c**) Deletion of long fragments containing two essential genes. (**d**) Schematic of the deletion of fragment No.1. LHA: left homologous arm. RHA: right homologous arm. F: forward primer. R: reverse primer. (**e**) Representative results of PCR verification in the deletion experiment of fragment No.1. (**f**) Results in the deletion experiments of 12 nonessential fragments. (**g**) Summary of cumulative deletion. Data are expressed as means ± s.d. from three independent experiments.

### 3.4 Identification of nonessential sequences and chromosomal simplification

According to previous reports, the MG1655 chromosome harbors 4497 genes, including 4296 protein-encoding genes and 201 RNA-encoding genes [38, 39]. Researchers at Keio University identified the essentiality of all protein-encoding genes in *E. coli* K-12 by single gene deletion, generating the Keio collection [40, 41]. This provided important information for us to identify potential nonessential long fragments in the MG1655 genome. To delete a long fragment, we needed to construct a plasmid#2 that expressed a pair of sgRNA targeting two flanks of the fragment and harboring the corresponding donor DNA (Fig. 4a). To delete a long fragment harboring a limited number of essential genes, we added these genes to the corresponding plasmid#2 between the two homologous arms. Therefore, the essential genes remained in the genome after editing, and the edited cells survived (Fig. 4b and 4c). For each long fragment deletion, we designed two pairs of primers for PCR verification. The first primer pair targets DNA sequences within the long fragment, and the second primer pair targets the adjacent sequences outside the two homologous arms (Fig. 4d and Fig. S6). The correct clones did not obtain PCR product using the first primer pair but obtained the corresponding PCR products using the second. On the contrary, the unedited control clone obtained the corresponding PCR products using the first primer pair but did not obtain PCR products using the second (Fig. 4e and Fig. S7).

Altogether, we successfully deleted 12 long nonessential fragments in the MG1655 genome (Table 1), including the 99.9-kb fragment (No. 3) mentioned in the previous section. These fragments are located in different regions of the genome, and their lengths range from 52.0 to 186.7 kb. Among the 12 fragments, No. 3, No. 8, and No. 11 harbor one essential gene; No. 1 and No. 4 harbor two essential genes; and No. 9 harbors three essential genes (Table 1). Based on the results of PCR and sequencing, we determined the editing efficiencies and positive rates (Fig. 4f). All positive rates were over 95%, and the editing efficiencies ranged from 2.3 × 10^−4^ to 1.3 × 10^−3^. The deletion of fragments No. 3, No. 4, and No. 7 led to much lower editing efficiencies than those from deletion of the other fragments. By measuring growth curves of the 12 knockout strains, we found that the No. 3, No. 4, and No. 7 knockout strains grew much slower than other knockout strains, and the No. 4 knockout strain grew slowest (Fig. S5). This phenomenon implies that these fragments were important for cell growth, and that the lower viability of edited cells might have led to the increase in editing efficiency. In addition, the different DNA loci and sgRNA sequences are also possible factors influencing the editing efficiency.

**Table 1.** Long fragments deleted in the MG1655 genome.

| Fragment No. | Starting site | End site | Length | Essential gene |
| --- | --- | --- | --- | --- |
| No. 1 | 240,056 | 426,771 | 186,715 | *yagG, hemB* |
| No. 2 | 499,529 | 552,955 | 53,426 | None |
| No. 3 | 565,456 | 665,088 | 99,932 | *entD* |
| No. 4 | 990,473 | 1,127,061 | 136,588 | *fabA, serT* |
| No. 5 | 1,449,596 | 1,549,490 | 99,894 | None |
| No. 6 | 1,549,491 | 1,647,484 | 97,993 | None |
| No. 7 | 2,349,152 | 2,430,141 | 80,989 | None |
| No. 8 | 2,442,420 | 2,517,306 | 74,886 | *argW* |
| No. 9 | 2,822,534 | 2,906,555 | 84,021 | *ispF, ispD, ftsB* |
| No. 10 | 3,610,719 | 3,689,415 | 78,696 | None |
| No. 11 | 3,824,765 | 3,876,879 | 52,114 | *selC* |
| No. 12 | 4,198,958 | 4,251,002 | 52,044 | None |

After deleting 12 long fragments individually, we tried to construct cumulative deletion mutants. Here, we used MG1655-ΔNo. X to represent the MG1655 mutant that loses fragment No. X (X=1, 2, 3, …, 12). As No. 1 was the longest fragment deleted in this study (Table 1), we chose to construct cumulative deletion mutants based on strain MG1655-ΔNo. 1. Though iterative editing, we successfully deleted fragment No. 9 from MG1655-ΔNo. 1, generating strain MG1655-ΔNo. 1/ΔNo. 9 that lost a total of 270.7 kb of the DNA sequence, containing 268 open reading frames (ORFs) (Fig. 4g). We then tried to delete a third fragment based on MG1655-ΔNo. 1/ΔNo. 9. According to the growth curves of single deletion mutants, the knockout of fragment No. 2, No. 5, No. 6, No. 8, No. 10, or No. 12 had no apparent influence on cell growth (Fig. S5). Therefore, we attempted to delete these fragments individually in MG1655-ΔNo. 1/ΔNo. 9. As a result, we successfully obtained strains MG1655-ΔNo. 1/ΔNo. 9/ΔNo. 2, MG1655-ΔNo. 1/ΔNo. 9/ΔNo. 5, and MG1655-ΔNo. 1/ΔNo. 9/ΔNo. 6. The three knockout strains lost a total of 324.1 kb, 370.6 kb, and 368.7 kb of the DNA sequences containing 315, 364, and 368 ORFs, respectively (Fig. 4g). We failed to knock out fragments No. 8, No. 10, and No. 12 in MG1655-ΔNo. 1/ΔNo. 9 despite repeating the experiments several times, implying that these fragments were all essential for the survival of MG1655-ΔNo. 1/ΔNo. 9.

### 3.5 Metabolic engineering of *E. coli* for isobutanol production

Higher alcohols such as isobutanol and n-butanol show promise in becoming the next generation of biofuels, due to their higher energy density, higher vapor pressure, and relatively low hydroscopicity [42, 43]. To illustrate the potential of applying the genome editing method to metabolic engineering, we used the method to modify the *E. coli* genome for producing isobutanol. First, we constructed a chassis strain named JW74 based on MG1655 with six rounds of genomic editing (Fig. 5a). The competency of JW74 was 170-fold that of MG1655, making it much easier to transform exogenous DNA. We then built a 7.9-kb operon and integrated it into the JW74 chromosome, thus displacing fragment No. 5 (Fig. 5a) and generating strain SH258. Fragment No. 5 was 99.9 kb in length, and the corresponding knockout strain grew slightly faster than its parental strain (Fig. S5f). The operon consists of five structural genes and 5′ and 3′ untranslated regions (UTRs). The 5′ UTR contains a strong bacterial ribosome-binding site [44] and a T7 promoter, which naturally controls the expression of bacteriophage T7 RNA polymerase [45]; the 3′ UTR contains a T7 terminator. The five structural genes are *alsS, ilvC, ilvD, kivD*, and *adhA* (Fig. 5a). Among the five genes, *ilvC* and *ilvD* came from *E. coli, alsS* came from *Bacillus subtilis* [46], and *kivD* and *adhA* came from *Lactococcus lactis* [47] (Fig. 5b). In order to initiate transcription of the operon, we introduced the T7 RNA polymerase-encoding gene controlled by the T5 promoter [48] to the SH258 genome, generating the SH274 strain (Fig. 5a). Though the T5 promoter is an inducible promoter repressed by LacI, it served here as a strong constitutive promoter. This is because SH274 is a *lacI*-defective strain. In traditional metabolic engineering, introducing a high-copy-number fermentation plasmid is a commonly used strategy to overexpress enzymes related to the target products. Therefore, we constructed the pColE1-P_T5_-*alsS*-*ilvC*-*ilvD*-*kivD*-*adhA* plasmid and transformed it into JW74, generating the SH279 strain.

**Fig.5.**
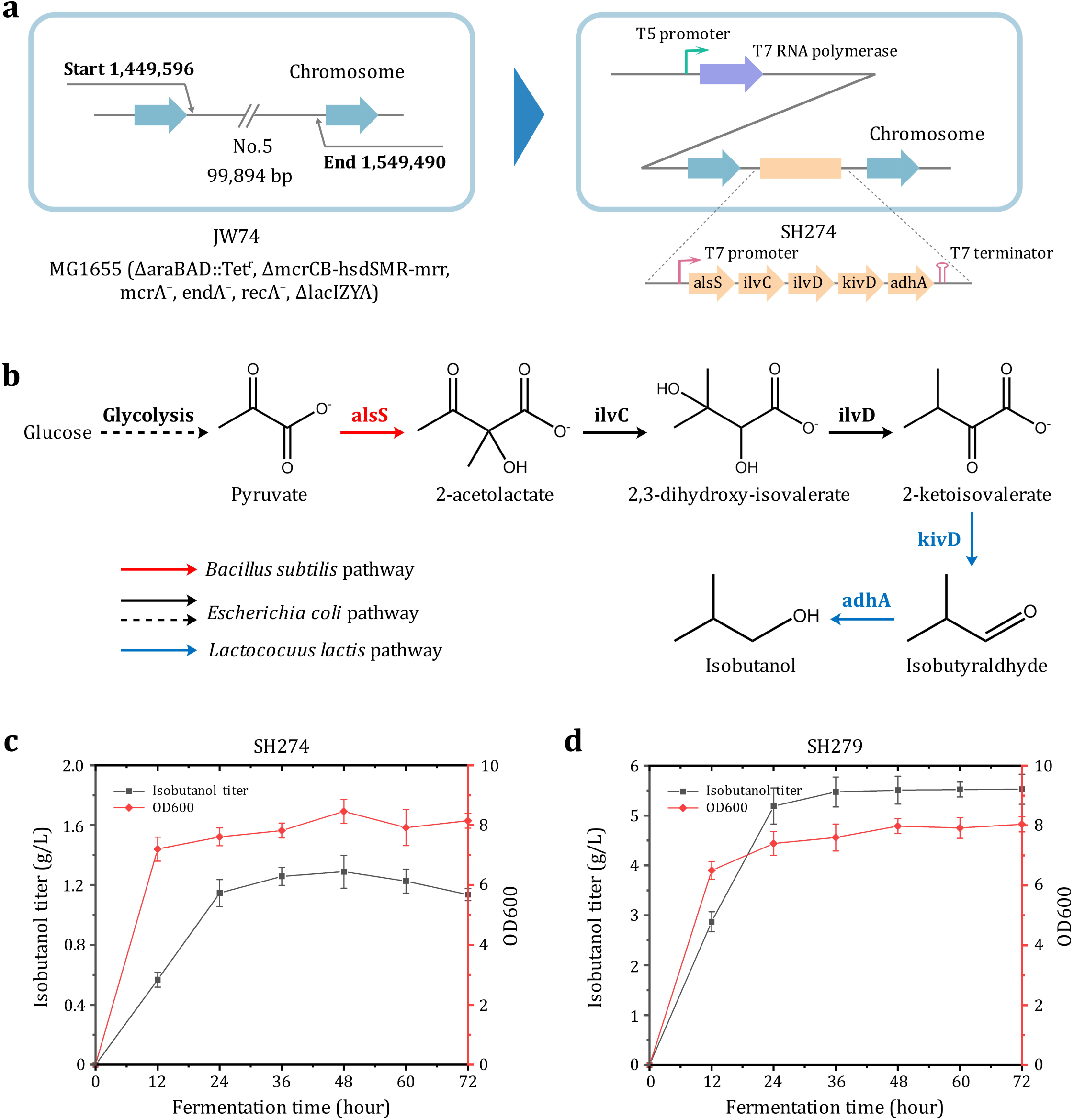
Metabolic engineering of *E. coli* for isobutanol production. (**a**) Construction of the SH274 strain based on the JW74 strain. (**b**) The synthetic pathway of isobutanol. (**c**) Results of isobutanol fermentation of the SH274 strain. (**d**) Results of isobutanol fermentation of the SH279 strain. Data are expressed as means ± s.d. from three independent experiments.

We used the strains SH274 and SH279 to conduct micro-aerobic fermentation in shake flasks containing 20 mL of M9 medium. Briefly, the acetolactate synthase (AlsS) converts pyruvate, the intermediate product of glycolysis, into 2-acetolactate. This is then transformed into 2,3-dihydroxy-isovalerate by ketol-acid reductoisomerase (IlvC). As the substrate of dihydroxyacid dehydratase (IlvD), 2,3-dihydroxy-isovalerate is converted into 2-ketoisovalerate, which is transformed into isobutyraldhyde by 2-ketoisovalerate decarboxylase (KivD). Finally, isobutyraldhyde is catalyzed by alcohol dehydrogenase (AdhA), generating isobutanol (Fig. 5b). During fermentation, samples were taken every 12 h to measure the OD600 value and isobutanol titer (Fig. 5c). As a result, isobutanol reached a maximum titer of 1.3 g/L after 48 h of SH274 fermentation (Fig. 5c). To our knowledge, this was the first attempt to produce isobutanol without introducing a high-copy-number fermentation plasmid, and isobutanol production was higher than many reports using such a plasmid [49, 50]. For strain SH279, isobutanol reached a maximum titer of 5.5 g/L after 48 h (Fig. 5d). This is 4.2-fold that of SH274, indicating that the SH274 strain has much room for improvement. In future study, we therefore plan to increase the copy number of the operon P_T7_-*alsS*-*ilvC*-*ilvD*-*kivD*-*adhA*-T_T7_ in the SH274 genome to strengthen the expression of related enzymes.

## 5 Discussion

Many CRISPR/Cas9-based methods have been developed for genome engineering in *E. coli*. In these methods, the editing efficiencies range from 10^−7^ to 10^−4^ [33]. Generally, the CRISPR/Cas9 system is combined with heterologous DSB repair systems, including homologous recombination systems [23-25, 27-29, 34, 35] or non-homologous end-joining (NHEJ) systems [36, 37]. NHEJ-mediated methods generate stochastic DNA indels in the target region, which makes genome editing inaccurate. In contrast, homologous recombination-mediated methods can achieve precise genome editing with higher editing efficiency. The method developed in this study is based on the CRISPR/Cas9-assisted recombineering, which is a class of methods that combine the CRISPR/Cas9 system and λ-Red system. One-plasmid [25], two-plasmid [24, 27, 34, 35], and three-plasmid [26] systems have been used in existing methods, and the two-plasmid system was adopted in our method. In the one-plasmid system, a huge plasmid needs to be constructed for each round of genome editing. In the three-plasmid system, two plasmids need to be constructed for each round of genome editing, and the plasmid curing is difficult. By comparison, the two-plasmid system is more convenient to use. CRISPR/Cas9-assisted recombineering methods need an artificial donor DNA as the editing template. Both cycle DNA (plasmid-borne dsDNA [24-26]) and linear DNA (PCR-amplified dsDNA [24, 27-29, 34, 35] or synthesized ssDNA [23, 27]) have been used as the editing template in existing methods. Editing template integrated into a plasmid can avoid the attack by DNA exonucleases and copy itself along with plasmid replication, which greatly increases homologous recombination efficiency, and thus increases editing efficiency. However, the homologous recombination between a broken genome and a plasmid generates either non-crossover product or crossover product (Fig. S8a). Therefore, there is a significant chance that the total plasmid will be integrated into the genome at the target site, leading to undesired recombination products (Fig. S8b). As the two kinds of recombination events are difficult to distinguish through conventional PCR verification method, we suspect that previous researchers have not noticed this phenomenon in their genome editing experiments. Although genome editing with linear editing templates circumvents the trouble caused by plasmid integration, it results in much lower editing efficiency due to the sensitivity of linear editing templates to DNA exonucleases. To solve this contradiction, we integrated the editing template into a plasmid of the two-plasmid system and added the target sequence to the two flanks of the editing template. Therefore, the editing template can copy itself along with plasmid replication and can be released from the plasmid by Cas9 cleavage. By using this strategy, the advantages of the cycle editing template and the linear editing template are combined in one method. The using of a strict inducible promoter to control the expression of Cas9 and sgRNA also improves the performance of the two-plasmid system. With this strategy, the transformants have enough time to recover and reproduce before DNA cleavage, which increases the cell activity and initial cell number.

The genome engineering method developed in this study was efficient for different kinds of genome editing, including sequence insertion, sequence deletion, and sequence displacement. The method was particularly advantageous for long fragment editing, resulting in high editing efficiency and positive rates. With the aid of this method, we were able to insert DNA fragments up to 12 kb into the genome and to delete DNA fragments up to 186.7 kb from the genome. In the 12-kb insertion experiment, the positive rate was 98.3% and the editing efficiency was 7.2 × 10^−4^. In the 186.7-kb deletion experiment, the positive rate was 97.7% and the editing efficiency was 1.2 × 10^−3^. Compared with existing CRISPR/Cas9-based methods, our method results in higher editing efficiency and positive rates in long fragment editing. In addition, the high performance of this method is independent of high-competency host strains, making the method applicable to more experiments. Using this method, only “3*N* + 1” days are needed for “*N*” rounds of editing. To our knowledge, the 12 kb fragment and the 186.7 kb fragment inserted and deleted in this study are the longest fragments manipulated in the *E. coli* genome using CRISPR/Cas9-based genome editing methods. At present, our method can manipulate two targets in a single round of genome editing. In future studies, we will upgrade the method to manipulate more targets at the same time.

As a powerful genome engineering tool, the method has great application potential. In this study, to demonstrate its potential, we have applied the method in genome simplification and metabolic engineering. *E. coli* has been the prominent prokaryotic organism in research laboratories since the origin of molecular biology, and is arguably the most completely characterized single-cell life form [51]. According to previous studies, different *E. coli* strains possess different genome sizes. For example, MG1655, an *E. coli* K-12 strain, has a 4.6-Mb genome that harbors 4497 genes, including 4296 protein-encoding genes and 201 RNA-encoding genes [38, 39]. Functional analyses have shown that *E. coli* cells grown under given conditions use only a small fraction of their genes [52]. As Koob et al. have proposed, deletion of genes that are nonessential under a given set of growth conditions could identify a minimized set of essential *E. coli* genes and DNA sequences [53]. In past decades, researchers have explored nonessential sequences and removed them from the *E. coli* genome individually or cumulatively, trying to construct a minimized genome [1, 2, 54, 55]. In their studies, the methods utilized to delete nonessential sequences are very complicated and time-consuming. To remove a long fragment from the genome, researchers have tried many recombination techniques both alone or in combination, including Flp/*FRT*, Cre/*loxP*, λ-Red, Tn5 transposon, and phage P1 transduction [1, 2, 54, 56]. Compared with these methods, our method is time-saving and easy to handle. Using this method, we have constructed 12 individual-deletion and four cumulative-deletion strains based on MG1655, with the simplest genome lacking a total of 370.6 kb of DNA sequence containing 364 ORFs. Although some of the deletions generated could coexist in a single strain, many deletions that were viable individually were not viable when combined with other deletions, which clearly indicates that some genes are not dispensable simultaneously, despite being dispensable individually. The genes belonging to this group may be those involved in alternative metabolic pathways. This observation also suggests that the number of essential genes is greater than estimated, and further illustrates the utility of our combinatorial-deletion approach for functional study of the *E. coli* genome.

Microorganisms are versatile living systems for achieving biosynthesis of valuable molecules contributing to chemical, energy, and pharmaceutical processes [57-61]. Plasmids have been commonly used for domesticating microbial materials to obtain desired cellular functions, due to the simplicity of genetic manipulation. Inspired by nature, antibiotics have been widely used to minimize phenotype variation of plasmid-containing microbes. However, the use of antibiotics may result in multidrug-resistant species by horizontal gene transfer, and metabolic burden leading to suboptimal production of target compounds [4]. The addition of antibiotics not only increases the cost, but also contaminates final products in industrial settings. Genome integration is a good alternative to plasmids and provides more stability for artificially introduced genetic information. The technique we developed is efficient for genome integration. In this study, we integrated the isobutanol synthetic pathway into a chassis strain derived from MG1655, generating a genetically stable metabolic engineering strain that produced 1.3 g/L isobutanol in a shake flask. As expected, productivity of this engineered strain was lower than the strain containing a high-copy-number fermentation plasmid, mainly due to the low expression of related enzymes. In future studies, we will endeavor to increase isobutanol production by integrating more copies of the isobutanol synthetic pathway into the genome.

## 6 Conclusions

Overall, this study developed an efficient genome engineering method for the insertion and deletion of long DNA fragments in the *E. coli* genome and demonstrated the tool’s potential in synthetic biology by applying it in genome simplification and metabolic engineering.

## Acknowledgements

This work was jointly supported by The National Key Research and Development Program of China (No. 2019YFA0904100, to Yi-Xin Huo) and The National Natural Science Foundation of China (No. 31961133014, to Yi-Xin Huo).

## Compliance with ethics guidelines

All authors (Chaoyong Huang, Liwei Guo, Jingge Wang, Ning Wang, Yi-Xin Huo) declare that they have no conflict of interest or financial conflicts to disclose.

## References

1. J. Kato, M. Hashimoto. Construction of consecutive deletions of the *Escherichia coli* chromosome. Mol. Syst. Biol., 2007, 3: 132

2. J. Kato, M. Hashimoto. Construction of long chromosomal deletion mutants of *Escherichia coli* and minimization of the genome. Methods Mol. Biol., 2008, 416: 279–293

3. S. L. Shipman, J. Nivala, J. D. Macklis, G. M. Chruch. CRISPR-Cas encoding of a digital movie into the genomes of a population of living bacteria. Nature, 2017, 547: 345–349

4. C. Mignon, R. Sodoyer, B. Werle. Antibiotic-free selection in biotherapeutics: now and forever. Pathogens, 2015, 4: 157–181

5. A. Baudin, O. Ozier-Kalogeropoulos, A. Denouel, F. Lacroute, C. Cullin. A simple and efficient method for direct gene deletion in *Saccharomyces cerevisiae.* Nucleic Acids Res., 1993, 21: 3329–3330

6. R. B. Wilson, D. Davis, A. P. Mitchell. Rapid hypothesis testing with *Candida albicans* through gene disruption with short homology regions. J. Bacteriol., 1999, 181: 1868–1874

7. C. B. Russell, D. S. Thaler, F. W. Dahlquist. Chromosomal transformation of *Escherichia coli recD* strains with linearized plasmids J. Bacteriol., 1989, 171: 2609–2613

8. A. J. Link, D. Phillips, G. M. Church. Methods for generating precise deletions and insertions in the genome of wild-type *Escherichia coli*: application to open reading frame characterization. J. Bacteriol., 1997, 179: 6228–6237

9. G. Pósfai, V. Kolisnychenko, Z. Bereczki, F. R. Blattner. Markerless gene replacement in *Escherichia coli* stimulated by a double-strand break in the chromosome. Nucleic Acids Res., 1999, 27: 4409–4415

10. S. K. Sharan, L. C. Thomason, S. G. Kuznetsov, D. L. Court. Recombineering: a homologous recombination-based method of genetic engineering. Nat. Protoc., 2009, 4: 206–223

11. J. Jeong, N. Cho, D. Jung, D. Bang. Genome-scale genetic engineering in *Escherichia coli.* Biotechnol. Adv., 2013, 31: 804–810

12. G. Pines, E. F. Freed, J. D. Winkler, R. T. Gill. Bacterial recombineering: genome engineering via phage-based homologous recombination. ACS Synth. Biol., 2015, 4: 1176–1185

13. K. A. Datsenko, B. L. Wanner. One-step inactivation of chromosomal genes in *Escherichia coli* K-12 using PCR products. Proc. Natl. Acad. Sci. U.S.A., 2000, 97: 6640–6645

14. H. Wang, X. Bian, L. Xia, X. Ding, R. Müller, Y. Zhang, et al. Improved seamless mutagenesis by recombineering using *ccdB* for counterselection. Nucleic Acids Res., 2014, 42: e37

15. H. H. Wang, F. J. Isaacs, P. A. Carr, Z. Z. Sun, G. Xu, C. R. Forest, et al. Programming cells by multiplex genome engineering and accelerated evolution. Nature, 2009, 460: 894–898

16. J. R. Warner, P. J. Reeder, A. Karimpour-Fard, L. B. Woodruff, R. T. Gill. Rapid profiling of a microbial genome using mixtures of barcoded oligonucleotides. Nat. Biotechnol., 2010, 28: 856–862

17. F. J. Isaacs, P. A. Carr, H. H. Wang, M. J. Lajoie, B. Sterling, L. Kraal, et al. Precise manipulation of chromosomes in vivo enables genome-wide codon replacement. Science, 2011, 333: 348–353

18. B. K. Tischer, J. von Einem, B. Kaufer, N. Osterrieder. Two-step Red-mediated recombination for versatile high-efficiency markerless DNA manipulation in *Escherichia coli.* Biotechniques, 2006, 40: 191–197

19. B. J. Yu, K. H. Kang, J. H. Lee, B. H. Sung, M. S. Kim, S. C. Kim. Rapid and efficient construction of markerless deletions in the *Escherichia coli* genome. Nucleic Acids Res., 2008, 36: e84

20. J. Yang, B. Sun, H. Huang, Y. Jiang, L. Diao, B. Chen, et al. High-efficiency scarless genetic modification in *Escherichia coli* by using lambda red recombination and I-SceI cleavage. Appl. Environ. Microbiol., 2014, 80: 3826–3834

21. M. Jinek, K. Chylinski, I. Fonfara, M. Hauer, J. A. Doudna, E. Charpentier. A programmable dual RNA-guided DNA endonuclease in adaptive bacterial immunity. Science, 2012, 337: 816–821

22. G. Gasiunas, R. Barrangou, P. Horvath, V. Siksnys. Cas9-crRNA ribonucleoprotein complex mediates specific DNA cleavage for adaptive immunity in bacteria. Proc. Natl. Acad. Sci. U.S.A., 2012, 109: e2579–2586

23. W. Jiang, D. Bikard, D. Cox, F. Zhang, L. A. Marraffini. RNA-guided editing of bacterial genomes using CRISPR-Cas systems. Nat. Biotechnol., 2013, 31: 233–239

24. Y. Jiang, B. Chen, C. Duan, B. Sun, J. Yang, S. Yang. Multigene editing in the *Escherichia coli* genome via the CRISPR-Cas9 system. Appl. Environ. Microbiol., 2015, 81: 2506

25. D. Zhao, S. Yuan, B. Xiong, H. Sun, L. Ye, J. Li, et al. Development of a fast and easy method for *Escherichia coli* genome editing with CRISPR/Cas9. Microb. Cell Fact., 2016, 15: 205

26. X. Feng, D. Zhao, X. Zhang, X. Ding, C. Bi. CRISPR/Cas9 assisted multiplex genome editing technique in *Escherichia coli*. Biotechnol. J., 2018, 13: 1700604

27. Y. Li, Z. Lin, C. Huang, Y. Zhang, Z. Wang, Y. Tang, et al. Metabolic engineering of *Escherichia coli* using CRISPR–Cas9 meditated genome editing. Metab. Eng., 2015, 31: 13–21

28. D. Zhao, X. Feng, X. Zhu, T. Wu, X. Zhang, C. Bi. CRISPR/Cas9-assisted gRNA-free one-step genome editing with no sequence limitations and improved targeting efficiency. Sci. Rep., 2017, 7: 16624

29. H. Zhang, Q. X. Cheng, A. M. Liu, G. P. Zhao, J. Wang. A novel and efficient method for bacteria genome editing employing both CRISPR/Cas9 and an antibiotic resistance cassette. Front. Microbiol., 2017, 8: 812

30. D. G. Gibson, L. Young, R. Y. Chuang, J. C. Venter, C. A. Hutchison, H. O. Smith. Enzymatic assembly of DNA molecules up to several hundred kilobases. Nat. Methods, 2009, 6: 343–345

31. M. G. Thompson, N. Sedaghatian, J. F. Barajas, M. Wehrs, C. B. Bailey, N. Kaplan, et al. Isolation and characterization of novel mutations in the pSC101 origin that increase copy number. Sci. Rep., 2018, 8: 1590

32. R. Chayot, B. Montagne, D. Mazel, M. Ricchetti. An end-joining repair mechanism in *Escherichia coli.* Proc. Natl. Acad. Sci. U.S.A., 2010, 107: 2141–2146

33. C. Huang, T. Ding, J. Wang, X. Wang, L. Guo, J. Wang, et al. CRISPR-Cas9-assisted native end-joining editing offers a simple strategy for efficient genetic engineering in *Escherichia coli.* Appl. Microbiol. Biotechnol., 2019, 103: 8497–8509

34. M. E. Chung, I. H. Yeh, L. Y. Sung, M. Y. Wu, Y. P. Chao, I. S. Ng, et al. Enhanced integration of large DNA into *E. coli* chromosome by CRISPR/Cas9. Biotechnol. Bioeng., 2017, 114: 172–183

35. Y. Li, F. Yan, H. Wu, G. Li, Y. Han, Q. Ma, et al. Multiple-step chromosomal integration of divided segments from a large DNA fragment via CRISPR/Cas9 in *Escherichia coli.* J. Ind. Microbiol. Biotechnol., 2019, 46: 81–90

36. T. Su, F. Liu, P. Gu, H. Jin, Y. Chang, Q. Wang, et al. A CRISPR-Cas9 assisted non-homologous end-joining strategy for one-step engineering of bacterial genome. Sci. Rep., 2016, 6: 37895

37. X. Zheng, S. Y. Li, G. P. Zhao, J. Wang. An efficient system for deletion of large DNA fragments in *Escherichia coli* via introduction of both Cas9 and the non-homologous end joining system from *Mycobacterium smegmatis.* Biochem. Biophys. Res., 2017, 485: 768–774

38. I. M. Keseler, A. Mackie, M. Peralta-Gil, A. Santos-Zavaleta, S. Gama-Castro, C. Bonavides-Martínez, et al. EcoCyc: fusing model organism databases with systems biology. Nucleic Acids Res., 2013, 41: D605–D612

39. I. M. Keseler, A. Mackie, A. Santos-Zavaleta, R. Billington, C. Bonavides-Martínez, R. Caspi, et al. The EcoCyc database: reflecting new knowledge about *Escherichia coli* K-12. Nucleic Acids Res., 2017, 45: D513–D550

40. N. Yamamoto, K. Nakahigashi, T. Nakamichi, M. Yoshino, Y. Takai, Y. Touda, et al. Update on the Keio collection of *Escherichia coli* single-gene deletion mutants. Mol. Syst. Biol., 2009, 5: 335

41. T. Baba, T. Ara, M. Hasegawa, Y. Takai, Y. Okumura, M. Baba, et al. Construction of *Escherichia coli* K-12 in-frame, single-gene knockout mutants: the Keio collection. Mol. Syst. Biol., 2006, 2: 8

42. M. Saini, Z. W. Wang, C. J. Chiang, Y. P. Chao. Metabolic engineering of *Escherichia coli* for production of n-butanol from crude glycerol. Biotechnol. Biofuels, 2017, 10: 173

43. S. Liang, H. Chen, J. Liu, J. Wen. Rational design of a synthetic Entner-Doudoroff pathway for enhancing glucose transformation to isobutanol in *Escherichia coli.* J. Ind. Microbiol. Biotechnol., 2018, 45: 187–199

44. M. B. Elowitz, S. Leibler. A synthetic oscillatory network of transcriptional regulators. Nature, 2000, 403: 335–338

45. M. Rong, B. He, W. T. McAllister, R. K. Durbin. Promoter specificity determinants of T7 RNALJpolymerase. Proc. Natl. Acad. Sci. U.S.A., 1998, 95: 515–519

46. S. Atsumi, Z. Li, J. C. Liao. Acetolactate synthase from *Bacillus subtilis* serves as a 2-ketoisovalerate decarboxylase for isobutanol biosynthesis in *Escherichia coli.* Appl. Environ. Microbiol., 2009, 75: 6306–6311

47. S. Atsumi, T. Hanai, J. C. Liao. Non-fermentative pathways for synthesis of branched-chain higher alcohols as biofuels. Nature, 2008, 451: 86–89

48. H. Bujard, R. Gentz, M. Lanzer, D. Stueber, M. Mueller, I. Ibrahimi, et al. A T5 promoter-based transcription-translation system for the analysis of proteins in vitro and in vivo. Methods Enzymol., 1987, 155: 416–433

49. C. T. Chen, J. C. Liao. Frontiers in microbial 1-butanol and isobutanol production. FEMS Microbiol. Lett., 2016, 363: fnw020

50. E. I. Lan, J. C. Liao. Microbial synthesis of n-butanol, isobutanol, and other higher alcohols from diverse resources. Bioresour. Technol., 2013, 135: 339–349

51. F. R. Blattner, G. Plunkett, C. A. Bloch, N. T. Perna, V. Burland, M. Riley, et al. The complete genome sequence of Escherichia coli K-12. Science, 1997, 277: 1453–1462

52. H. Tao, C. Bausch, C. Richmond, F. R. Blattner, T. Conway. Functional genomics: expression analysis of *Escherichia coli* growing on minimal and rich media. J. Bacteriol., 1999, 181: 6425–6440

53. M. D. Koob, A. J. Shaw, D. C. Cameron. Minimizing the Genome of *Escherichia coli.* Ann. N. Y. Acad. Sci., 1994, 745: 1–3

54. B. J. Yu, B. H. Sung, M. D. Koob, C. H. Lee, J. H. Lee, W. S. Lee, et al. Minimization of the *Escherichia coli* genome using a Tn5-targeted Cre/loxP excision system. Nat. Biotechnol., 2002, 20: 1018–1023

55. M. Hashimoto, T. Ichimura, H. Mizoguchi, K. Tanaka, K. Fujimitsu, K. Keyamura, et al. Cell size and nucleoid organization of engineered *Escherichia coli* cells with a reduced genome. Mol. Microbiol., 2005, 55: 137–149

56. Y. Kang, T. Durfee, J. D. Glasner, Y. Qiu, D. Frisch, K. M. Winterberg, et al. Systematic mutagenesis of the *Escherichia coli* genome. J. Bacteriol., 2004, 186: 4921–4930

57. C. J. Paddon, J. D. Keasling. Semi-synthetic artemisinin: a model for the use of synthetic biology in pharmaceutical development. Nat. Rev. Microbiol., 2014, 12: 355–367

58. H. Fang, D. Li, J. Kang, P. Jiang, J. Sun, D. Zhang. Metabolic engineering of *Escherichia coli* for de novo biosynthesis of vitamin B_12_. Nat. Commun., 2018, 9: 4917

59. Y. Yu, X. Zhu, H. Xu, X. Zhang. Construction of an energy-conserving glycerol utilization pathways for improving anaerobic succinate production in *Escherichia coli.* Metab. Eng., 2019, 56: 181–189

60. Y. X. Huo, H. Ren, H. Yu, L. Zhao, S. Yu, Y. Yan, et al. CipA-mediating enzyme self-assembly to enhance the biosynthesis of pyrogallol in *Escherichia coli.* Appl. Microbiol. Biotechnol., 2018, 102: 10005–10015

61. Y. X. Huo, K. M. Cho, J. G. Rivera, E. Monte, C. R. Shen, Y. Yan, et al. Conversion of proteins into biofuels by engineering nitrogen flux. Nat. Biotechnol., 2011, 29: 346–351

