## Supplementary Data for "Efficient long fragment editing technique enables large-scale and scarless bacterial genome engineering"

**Supplementary Figure 1**

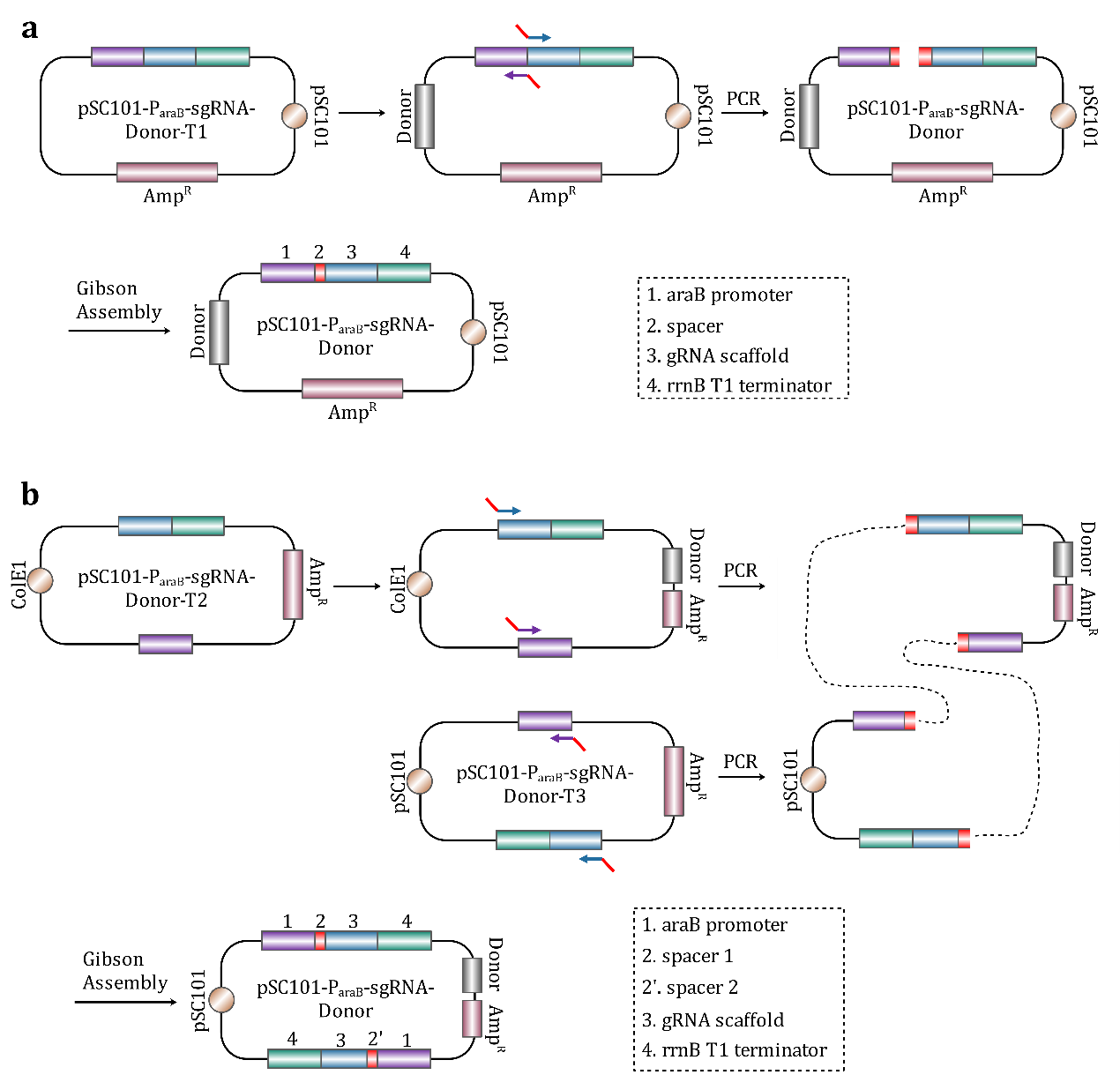

**Figure S1.** Construction procedures of plasmid pSC101-P_araB_-sgRNA-Donor. (**a**) Construction of plasmid pSC101-P_araB_-sgRNA-Donor containing one sgRNA expression chimera. The plasmid pSC101-P_araB_-sgRNA-Donor-T1 served as a parental plasmid, and a specifically designed donor DNA was integrated into it to construct an intermediate plasmid. The donor DNA contained two homologous arms that were about 500 bp. Then, a specific spacer (20 bp) was inserted into it between the araB promoter and the gRNA scaffold via single PCR and single Gibson Assembly. The spacer introduced by PCR served as overlap in Gibson Assembly. (**b**) Construction of plasmid pSC101-P_araB_-sgRNA-Donor containing two sgRNA expression chimeras. The plasmids pSC101-P_araB_-sgRNA-Donor-T2 and pSC101-P_araB_-sgRNA-Donor-T3 served as parental plasmids. First, a specifically designed donor DNA was integrated into plasmid pSC101-P_araB_-sgRNA-Donor-T2 to construct an intermediate plasmid. Then, the intermediate plasmid and the plasmid pSC101-P_araB_-sgRNA-Donor-T3 were combined to construct the plasmid pSC101-P_araB_-sgRNA-Donor through PCR and Gibson Assembly. The two specific spacers introduced by PCR served as overlaps in Gibson Assembly.

**Supplementary Figure 2**

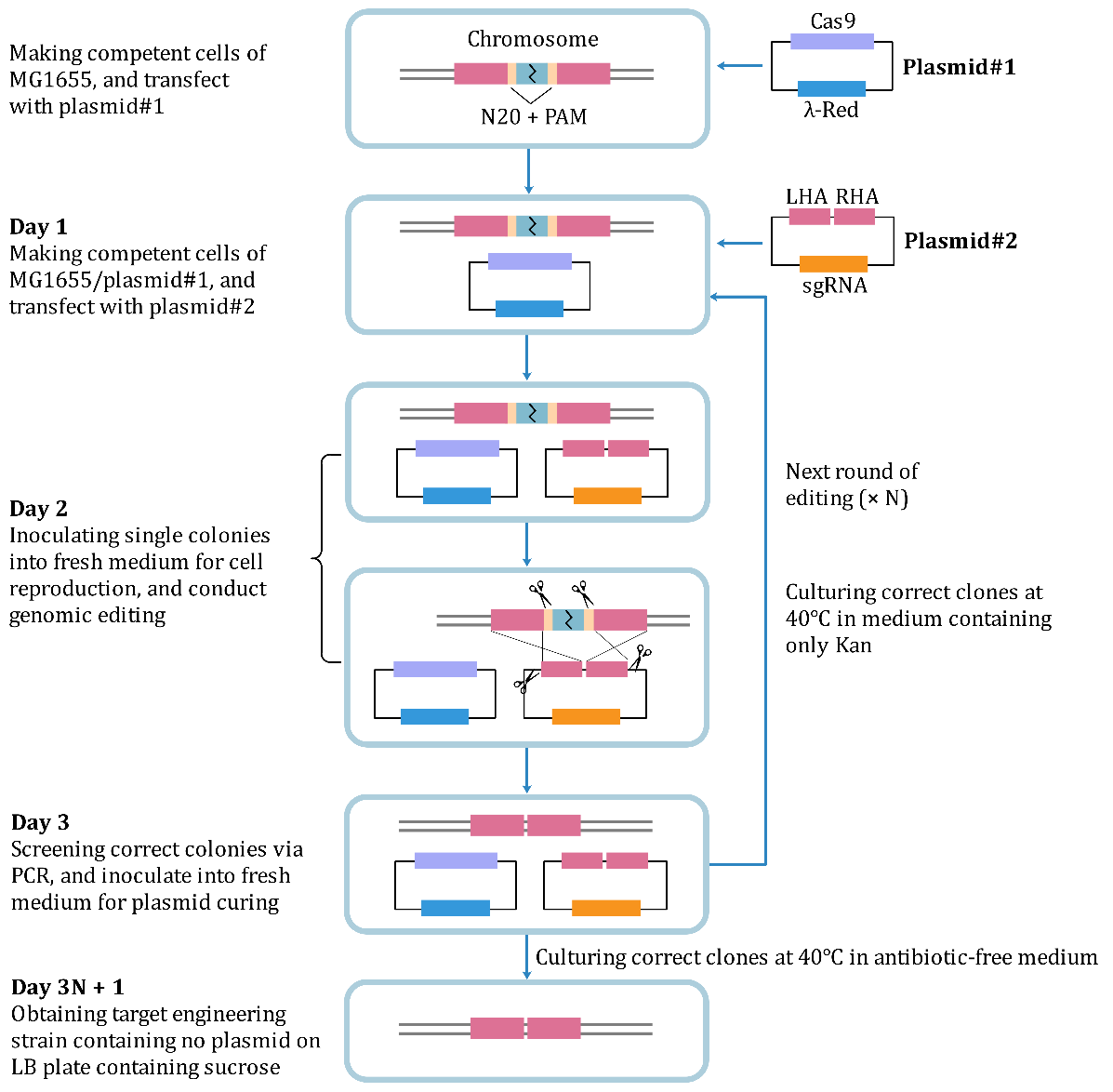

**Figure S2.** Flowchart of iterative editing. First, the Kan^R^ plasmid#1 was transfected into the target strain, such as MG1655, to obtain corresponding transformants, such as MG1655/plasmid#1. Then, specific plasmid#2 was transfected into the MG1655/plasmid#1 strain, and the MG1655/plasmid#1/plasmid#2 strain was screened in a LB plate with Amp, Kan and glucose at 30 °C. One or several single colonies were inoculated into 2 mL LB medium, and the culture was cultivated at 30°C for two hours. Then, 2 μL Amp, 2 μL Kan and 20 μL IPTG were added to the culture. After one hour, 20 μL L-arabinose was added, and the cultures were cultured for another three hours before plating. A 1-μL or 0.1-μL aliquot of the culture were plated onto a LB plate containing Amp, Kan and L-arabinose, and the plate was cultivated overnight at 30°C. Positive mutants were verified by colony PCR and sequencing. Positive mutant was cultivated in LB medium in the presence of only Kan at 40 °C for 12 hours to remove the temperature-sensitive Amp^R^ plasmid#2. Then, the obtained edited strain containing only the plasmid#1 was used as the starting strain for the next round of genomic editing. When the last round of genomic editing was completed, the edited strain was cultivated in LB medium without Kan at 40 °C for 12 hours to remove both Amp^R^ plasmid#2 and sucrose-sensitive Kan^R^ plasmid#1. The overnight culture was diluted for plating on a LB plate containing sucrose. Theoretically, colonies grow on the plate are plasmid-free.

**Supplementary Figure 3**

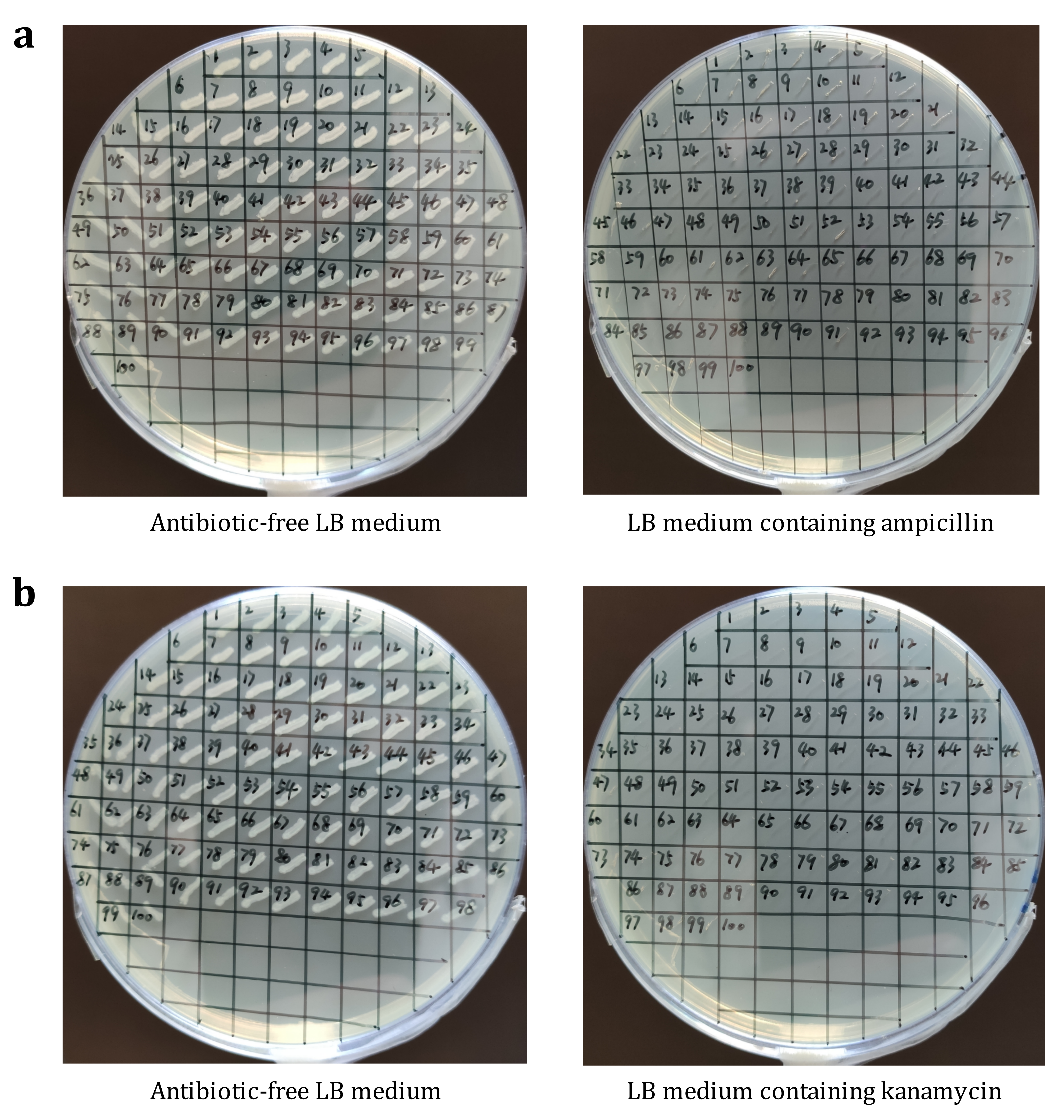

**Figure S3.** Results of plasmid curing. (**a**) Plasmid curing of plasmid#2. A single colony was inoculated into LB medium containing only Kan. Cells were cultivated at 40 °C for different time before plating on LB plates containing Kan. One hundred colonies were spotted on both antibiotic-free LB plate and LB plate containing Amp to test the loss rate of plasmid#2. (**b**) Plasmid curing of plasmid#1. A single colony was inoculated into LB medium containing no antibiotic. Cells were cultivated at 37 °C for different time before plating on antibiotic-free plate. One hundred colonies were spotted on both antibiotic-free LB plate and LB plate containing Kan to test the loss rate of plasmid#1.

**Supplementary Figure 4**

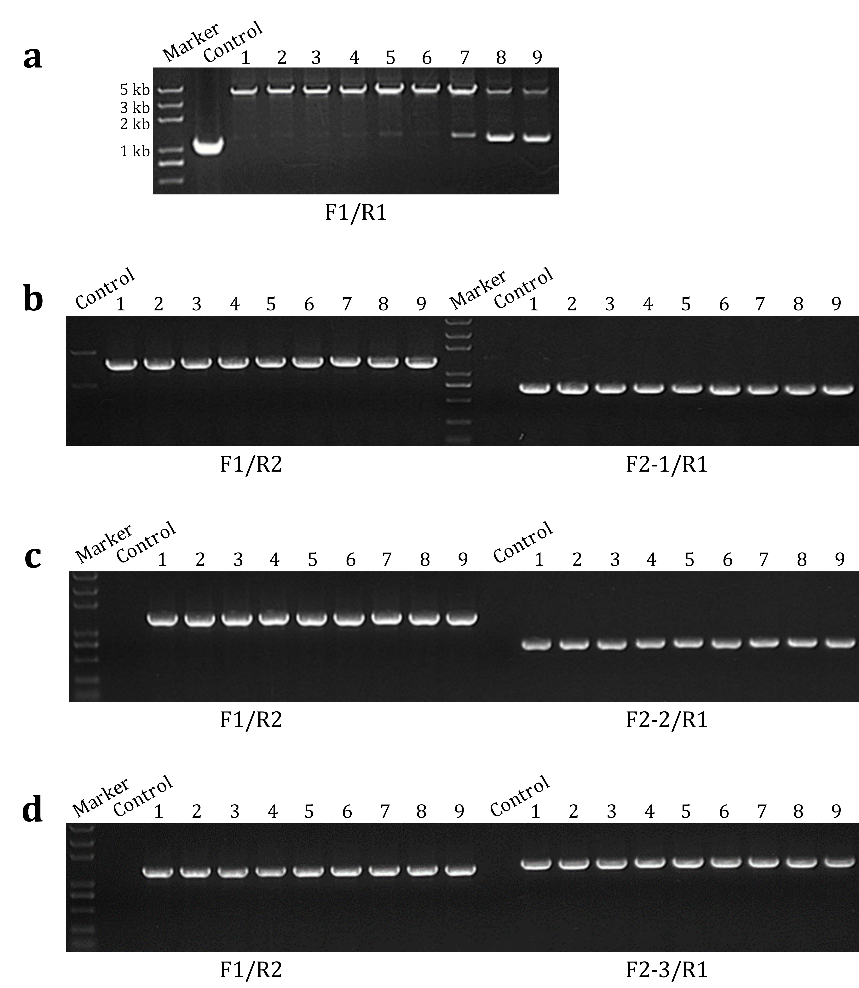

**Figure S4.** Representative results of PCR verification in large fragments insertion experiments. (**a**) PCR verification of 3-kb insertion. (**b**) PCR verification of 6-kb insertion. (**c**) PCR verification of 9-kb insertion. (**d**) PCR verification of 12-kb insertion.

**Supplementary Figure 5**

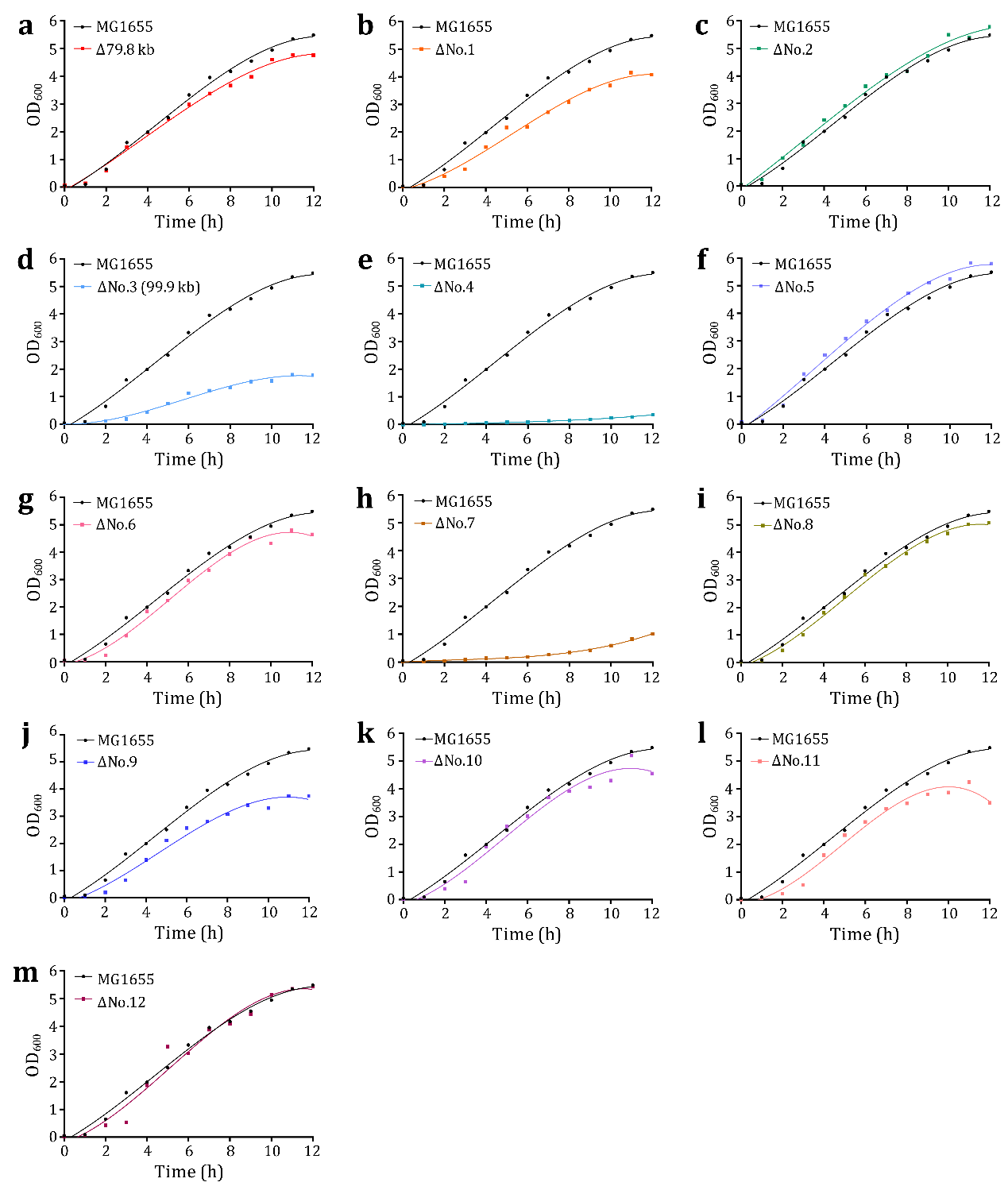

**Figure S5.** Comparison of growth curves between wild-type MG1655 and large fragment knockout strains. (**a**) Growth curve of strain MG1655-Δ79.8 kb. (**b**) Growth curve of strain MG1655-ΔNo.1. (**c**) Growth curve of strain MG1655-ΔNo.2. (**d**) Growth curve of strain MG1655-ΔNo.3. (**e**) Growth curve of strain MG1655-ΔNo.4. (**f**) Growth curve of strain MG1655-ΔNo.5. (**g**) Growth curve of strain MG1655-ΔNo.6. (**h**) Growth curve of strain MG1655-ΔNo.7. (**i**) Growth curve of strain MG1655-ΔNo.8. (**j**) Growth curve of strain MG1655-ΔNo.9. (**k**) Growth curve of strain MG1655-ΔNo.10. (**l**) Growth curve of strain MG1655-ΔNo.11. (**m**) Growth curve of strain MG1655-ΔNo.12.

**Supplementary Figure 6**

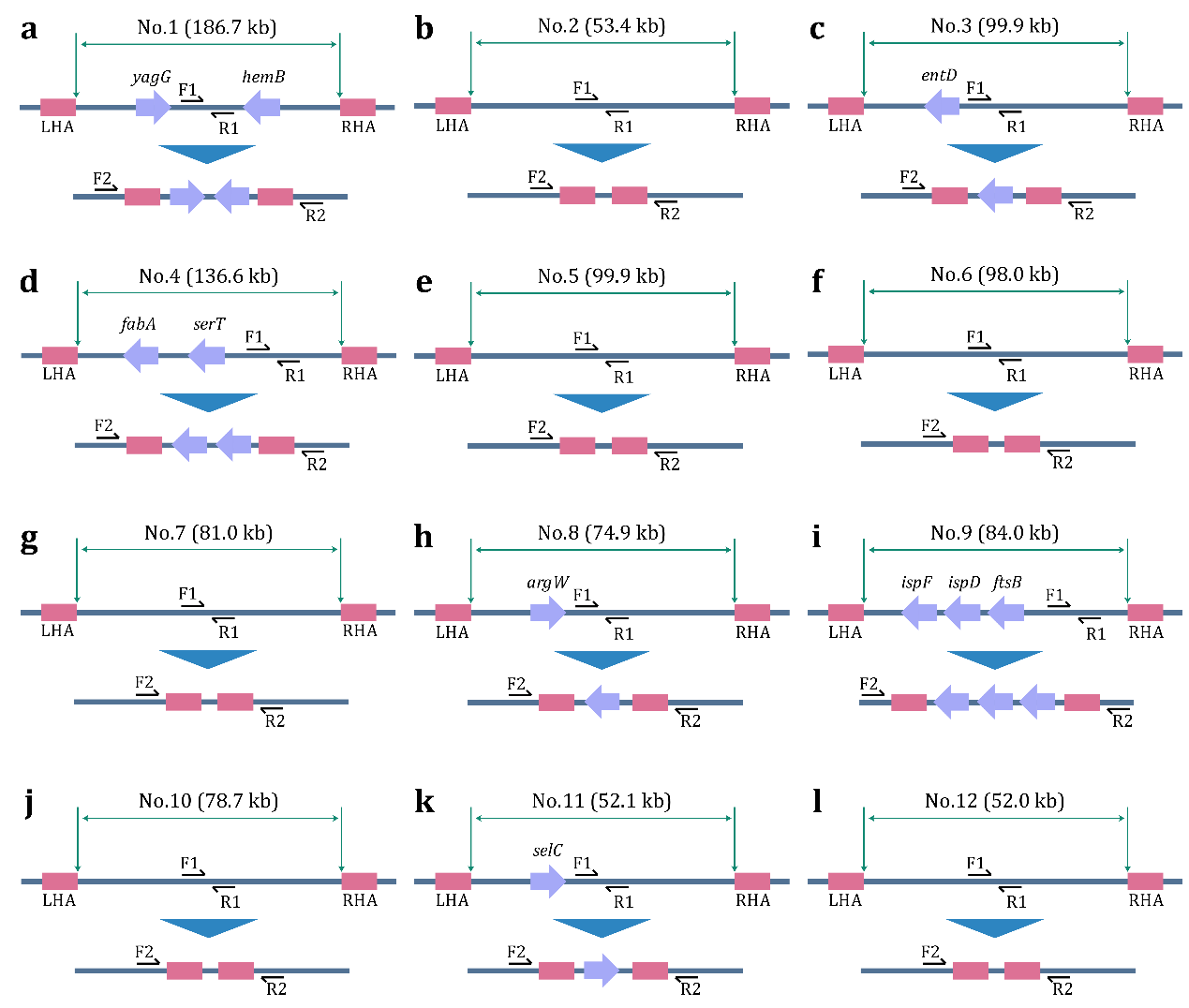

**Figure S6.** Schematic of the deletion of different long fragments. (**a**) Deletion of fragment No.1. (**b**) Deletion of fragment No.2. (**c**) Deletion of fragment No.3. (**d**) Deletion of fragment No.4. (**e**) Deletion of fragment No.5. (**f**) Deletion of fragment No.6. (**g**) Deletion of fragment No.7. (**h**) Deletion of fragment No.8. (**i**) Deletion of fragment No.9. (**j**) Deletion of fragment No.10. (**k**) Deletion of fragment No.11. (**l**) Deletion of fragment No.12.

**Supplementary Figure 7**

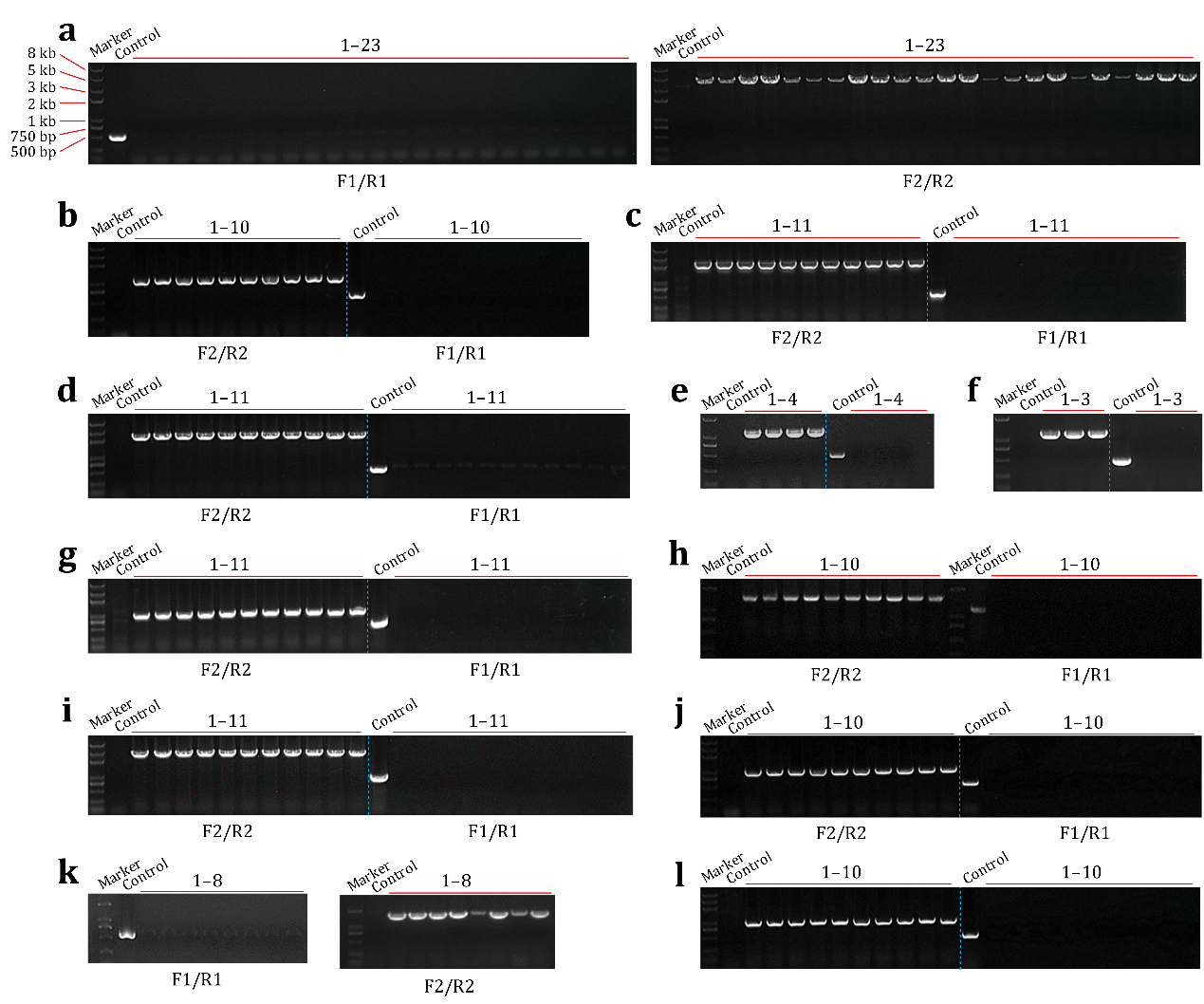

**Figure S7.** Representative results of PCR verification in large fragments deletion experiments. (**a**) PCR verification of fragment No.1 deletion. (**b**) PCR verification of fragment No.2 deletion. (**c**) PCR verification of fragment No.3 deletion. (**d**) PCR verification of fragment No.4 deletion. (**e**) PCR verification of fragment No.5 deletion. (**f**) PCR verification of fragment No.6 deletion. (**g**) PCR verification of fragment No.7 deletion. (**h**) PCR verification of fragment No.8 deletion. (**i**) PCR verification of fragment No.9 deletion. (**j**) PCR verification of fragment No.10 deletion. (**k**) PCR verification of fragment No.11 deletion. (**l**) PCR verification of fragment No.12 deletion.

**Supplementary Figure 8**

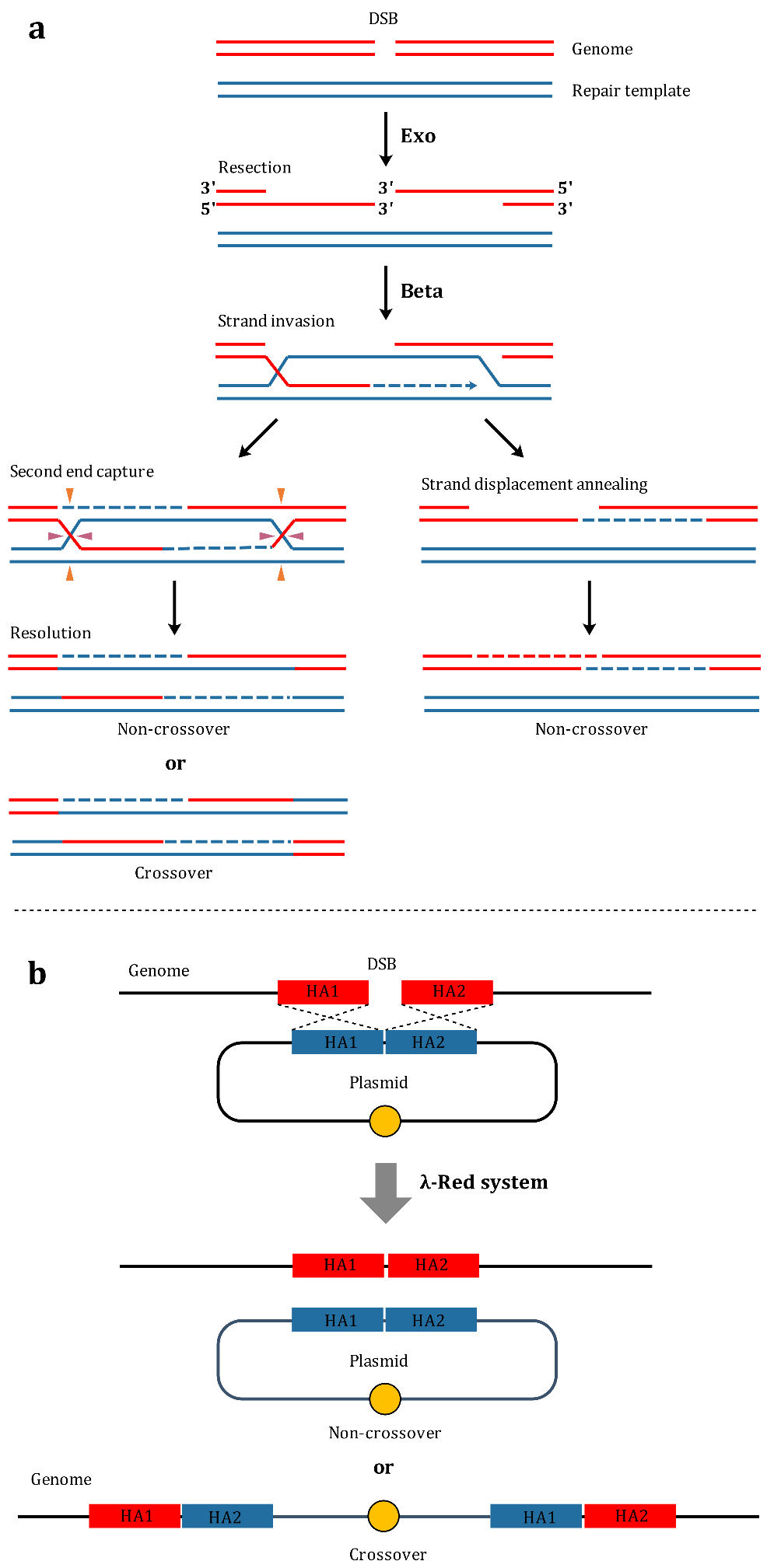

**Figure S8.** λ-Red-mediated homologous recombination. (**a**) Mechanism of λ-Red-mediated homologous recombination. (**b**) λ-Red-mediated homologous recombination between broken genome and plasmid.

**Table S1. Strains involved in this study**

| Strain | Description | Source/reference |
| --- | --- | --- |
| DH5α | F^–^ λ^–^ *endA1* *glnV44* *thi-1* *recA1* *relA1* *gyrA96* *deoR* *nupG* *purB20* φ80d*lacZ*ΔM15 Δ(*lacZYA*-*argF*)U169, hsdR17(*r_K_*^–^*m_K_*^+^) | [1] |
| MG1655 | F^–^ λ^–^ *ilvG*^–^ *rfb-50* *rph*-*1* | [2] |
| MG1655-*lacZ*::3kb | A 3-kb exogenous fragment was inserted into the *lacZ* gene | This study |
| MG1655-*lacZ*::6kb | A 6-kb exogenous fragment was inserted into the *lacZ* gene | This study |
| MG1655-*lacZ*::9kb | A 9-kb exogenous fragment was inserted into the *lacZ* gene | This study |
| MG1655-*lacZ*::12kb | A 12-kb exogenous fragment was inserted into the *lacZ* gene | This study |
| MG1655-Δ9.1kb | A 9.1-kb fragment (565,156–574,260) was deleted from the genome | This study |
| MG1655-Δ21.5kb | A 21.5-kb fragment (565,156–586,650) was deleted from the genome | This study |
| MG1655-Δ30.6kb | A 30.6-kb fragment (565,156–595,752) was deleted from the genome | This study |
| MG1655-Δ39.4kb | A 39.4-kb fragment (565,156–604,564) was deleted from the genome | This study |
| MG1655-Δ59.8kb | A 59.8-kb fragment (565,156–624,919) was deleted from the genome | This study |
| MG1655-Δ79.8kb | A 79.8-kb fragment (565,156–644,949) was deleted from the genome | This study |
| MG1655-Δ99.9kb (MG1655-ΔNo.3) | A 99.9-kb fragment (565,156–665,089) was deleted from the genome | This study |
| MG1655-ΔNo.1 | The fragment No.1 (240,056–426,771) was deleted from the genome | This study |
| MG1655-ΔNo.2 | The fragment No.2 (499,529–552,955) was deleted from the genome | This study |
| MG1655-ΔNo.4 | The fragment No.4 (990,473–1,127,061) was deleted from the genome | This study |
| MG1655-ΔNo.5 | The fragment No.5 (1,449,596–1,549,490) was deleted from the genome | This study |
| MG1655-ΔNo.6 | The fragment No.6 (1,549,491–1,647,484) was deleted from the genome | This study |
| MG1655-ΔNo.7 | The fragment No.7 (2,349,152–2,430,141) was deleted from the genome | This study |
| MG1655-ΔNo.8 | The fragment No.8 (2,442,420–2,517,306) was deleted from the genome | This study |
| MG1655-ΔNo.9 | The fragment No.9 (2,822,534–2,906,555) was deleted from the genome | This study |
| MG1655-ΔNo.10 | The fragment No.10 (3,610,719–3,689,415) was deleted from the genome | This study |
| MG1655-ΔNo.11 | The fragment No.11 (3,824,765–3,876,879) was deleted from the genome | This study |
| MG1655-ΔNo.12 | The fragment No.12 (4,198,958–4,251,002) was deleted from the genome | This study |
| MG1655-ΔNo.1/ΔNo.9 | The fragments No.1 and No.9 were deleted from the genome | This study |
| MG1655-ΔNo.1/ΔNo.9/ΔNo.2 | The fragments No.1, No.9 and No.3 were deleted from the genome | This study |
| MG1655-ΔNo.1/ΔNo.9/ΔNo.6 | The fragments No.1, No.9 and No.6 were deleted from the genome | This study |
| MG1655-ΔNo.1/ΔNo.9/ΔNo.5 | The fragments No.1, No.9 and No.5 were deleted from the genome | This study |
| JW74 | MG1655 (Δ*araBAD*::Tet^r^, Δ*mcrCB*-*hsdSMR*-*mrr*, *mcrA*^–^, *endA*^–^, *recA*^–^, Δ*lacIZYA*) | This study |
| SH258 | JW74 (Δfragment No.5::P_T7_-*alsS*-*ilvC*-*ilvD*-*kivD*-*adhA*-T_T7_) | This study |
| SH274 | JW74 (Δfragment No.5::P_T7_-*alsS*-*ilvC*-*ilvD*-*kivD*-*adhA*-T_T7_, P_T5_-RNAP (T7)) |  |
| SH279 | JW74 (pColE1-P_T5_-*alsS*-*ilvC*-*ilvD*-*kivD*-*adhA*) | This study |

**Table S2. Verification primers used in genomic editing experiments**

| Forward primer | Sequence (5’-3’) | Reverse primer | Sequence (5’-3’) | PCR product |
| --- | --- | --- | --- | --- |
| 3kb-F | CTGACTGGCGGTTAAATTGC | 3kb-R | TAACCGTCACGAGCATCATC | 4237 bp |
| 6kb-F1 | CTGACTGGCGGTTAAATTGC | 6kb-R1 | GGCCAGTTCGTTAAAATGCT | 1411 bp |
| 6kb-F2 | ACGCTGGACACACACATTAC | 6kb-R2 | TAACCGTCACGAGCATCATC | 744 bp |
| 9kb-F1 | CTGACTGGCGGTTAAATTGC | 9kb-R1 | GGCCAGTTCGTTAAAATGCT | 1411 bp |
| 9kb-F2 | TGCCCCATTTCACGACGTC | 9kb-R2 | TAACCGTCACGAGCATCATC | 754 bp |
| 12kb-F1 | CTGACTGGCGGTTAAATTGC | 12kb-R1 | GGCCAGTTCGTTAAAATGCT | 1411 bp |
| 12kb-F2 | AATCGTTCTCCCTGTATCGC | 12kb-R2 | TAACCGTCACGAGCATCATC | 1539 bp |
| Δ9.1kb-F1 | TTACATCGATGTGACAGGCC | Δ9.1kb-R1 | AAAACTCGCTTGTGGGAGCA | None |
| Δ9.1kb-F2 | TTCTTGATTCAGACGCGCAG | Δ9.1kb-R2 | GGGGATAACGCCTTAAATGG | 2065 bp |
| Δ21.5kb-F1 | GTGCTTCTAAAGGAAGTGGC | Δ21.5kb-R1 | AGAACCTTGAAACAGCATCC | None |
| Δ21.5kb-F2 | CGTTTGTTTGCCGCTATAGC | Δ21.5kb-R2 | TAAACTGACCGCTGAACAGC | 2251 bp |
| Δ30.6kb-F1 | CAATAAGGCGTTTTGCTTCC | Δ30.6kb-R1 | ATCTTGCCATGATAAAGGCC | None |
| Δ30.6kb-F2 | TTTTCTTGATTCAGACGCGC | Δ30.6kb-R2 | AATTTAACGCTTTGCAGGCT | 2285 bp |
| Δ39.4kb-F1 | AGCCTGAAAAAAGTATCGGG | Δ39.4kb-R1 | CTCACTACAACGCACCTGAA | None |
| Δ39.4kb-F2 | AATCAGAAAATCAGACGCGG | Δ39.4kb-R2 | AGCGTAAAATGCTTGATGCC | 2187 bp |
| Δ59.8kb-F1 | GCCAGTAACGAAAATCACCC | Δ59.8kb-R1 | GAGTTTTCAGAGCTGCATGG | None |
| Δ59.8kb-F2 | AATCAGAAAATCAGACGCGG | Δ59.8kb-R2 | TTTTTCTGACGCCAGCAGAC | 2217 bp |
| Δ79.8kb-F1 | AAGGTGGGCGCTATATCAAC | Δ79.8kb-R1 | CCAGATACAACAGCACGAAG | None |
| Δ79.8kb-F2 | TTCTTGATTCAGACGCGCAG | Δ79.8kb-R2 | ATTAATGCTACTGCGGTTGG | 2304 bp |
| Δ99.9kb-F1 | GCTTGCCGTCAATCAAAATC | Δ99.9kb-R1 | CCTACTTTTAACGCCGTCAC | None |
| Δ99.9kb-F2 | TTCTTGATTCAGACGCGCAG | Δ99.9kb-R2 | TGATGCATGATTACCAGCGC | 2214 bp |
| ΔNo.1-F1 | GCACACTGCGAAAAGATTGT | ΔNo.1-R1 | GCGCCAAAAGCTCTTTTACA | None |
| ΔNo.1-F2 | CCCACTCACCACAACCTAAAC | ΔNo.1-R2 | CCGAATACCACCAGCATCAA | 4265 bp |
| ΔNo.2-F1 | AAACCTAATTTTTTGACGACTTCATCG | ΔNo.2-R1 | CGGCGAAGCAACAACATAAAC | None |
| ΔNo.2-F2 | GAGCGGTAAAGAACTGAGCG | ΔNo.2-R2 | TGCTCGAACTTTATGGTCGC | 1188 bp |
| ΔNo.4-F1 | GACACATTGCCGACCAAACA | ΔNo.4-R1 | CTCTGGAAGCATGGTTGCAT | None |
| ΔNo.4-F2 | GATGGAAGCGCGACATATCA | ΔNo.4-R2 | TATGGTGAATTGCTACCGCC | 2468 bp |
| ΔNo.5-F1 | CGGTCGATATGCGGATGTAT | ΔNo.5-R1 | TCTCCTGCTGCAGGATTTTG | None |
| ΔNo.5-F2 | GTCGGTGTGTGTACGGTATT | ΔNo.5-R2 | ACATACGGGAACTGCTCTTT | 1181 bp |
| ΔNo.6-F1 | GACAACAAGCCCCTGATTAC | ΔNo.6-R1 | GAACTGGGGAGGCGACTATT | None |
| ΔNo.6-F2 | GGAGCAACTATCCGAAGTTC | ΔNo.6-R2 | CATGTTGGTGAGCATTGCAA | 1241 bp |
| ΔNo.7-F1 | GCATCCAGCGAATCAATTGC | ΔNo.7-R1 | GCGGCTAAAGCAAAAGCTAC | None |
| ΔNo.7-F2 | TTGAGTACCAGTGTCGCGAAG | ΔNo.7-R2 | GGGTTGTGCTTTCTTTGTTGC | 1181 bp |
| ΔNo.8-F1 | GTCGGCAAATTAACCGCATA | ΔNo.8-R1 | GCCAACCAGCACCATAATGA | None |
| ΔNo.8-F2 | CCTGCCAACCTCGAATTTAAT | ΔNo.8-R2 | CCCGATCAGTGAGAGTAGTG | 1521 bp |
| ΔNo.9-F1 | GCTATCCAGCCACTTAACG | ΔNo.9-R1 | AACTGAGCATTGACGGCATGA | None |
| ΔNo.9-F2 | CCCTTACTATGGAAAAAGTCGAG | ΔNo.9-R2 | GAAGAATTCACTCACTTCCTGG | 2946 bp |
| ΔNo.10-F1 | GTTTCGCCAGACTCTTCCATTAG | ΔNo.10-R1 | CACGCAGTTGTTCCACTTTG | None |
| ΔNo.10-F2 | GTCACGCTGTTTACCAAAGGC | ΔNo.10-R2 | GGCCAGAACGGATTTCAATATCATTG | 1091 bp |
| ΔNo.11-F1 | ATTCGTTACTGGCATGTTGATAC | ΔNo.11-R1 | AGTTACTCCACCATCACCAAAT | None |
| ΔNo.11-F2 | TATTCCTGGCGACCCGATTA | ΔNo.11-R2 | GGTCAACACGTAACGCTTTAC | 1622 bp |
| ΔNo.12-F1 | GTTTCGCCAGACTCTTCCATTAG | ΔNo.12-R1 | CACGCAGTTGTTCCACTTTG | None |
| ΔNo.12-F2 | GTCACGCTGTTTACCAAAGGC | ΔNo.12-R2 | GGCCAGAACGGATTTCAATATCATTG | 1104 bp |
| ΔaraBAD::Tet^r^-F1 | GTCGCATCAGGCGTTACATA | ΔaraBAD::Tet^r^-R1 | GATCGCCGCAGAAGAGAAAC | None |
| ΔaraBAD::Tet^r^-F2 | TCAATCTGCCCGGCAAACA | ΔaraBAD::Tet^r^-R2 | CTGGCGGGAACAGCAAAATA | 2528 bp |
| ΔmcrCB-hsdSMR-mrr-F1 | CGGGTTAGTGGCGACAATAT | ΔmcrCB-hsdSMR-mrr-R1 | TAACCGCACTGGTCAGCAA | None |
| ΔmcrCB-hsdSMR-mrr-F2 | GGATCAATTCTGAGACCGCT | ΔmcrCB-hsdSMR-mrr-R2 | GATCTGGGCCTGCTGTTCAA | 1272 bp |
| mcrA^–^-F1 | AGACAAGATAAGCGGCACAA | mcrA^–^-R1 | CGACTCTCGATAGAACCACT | 641 bp |
| mcrA^–^-F2 | AGTGGTTCTATCGAGAGTCG | mcrA^–^-R2 | CACCTGAAGACAGGGGAATT | 674 bp |
| mcrA^–^-F3 | AGACAAGATAAGCGGCACAA | mcrA^–^-R3 | CACCTGAAGACAGGGGAATT | 1295 bp |
| endA^–^-F1 | CTATCCTCTGCCGCTATGAA | endA^–^-R1 | GCAACACAGAAGGGGTACAT | 652 bp |
| endA^–^-F2 | ATGTACCCCTTCTGTGTTGC | endA^–^-R2 | CGTTGCACATACGGGTTATG | 678 bp |
| endA^–^-F3 | CTATCCTCTGCCGCTATGAA | endA^–^-R3 | CGTTGCACATACGGGTTATG | 1310 bp |
| recA^–^-F1 | GCGTAGAATTTCAGCGCGTTA | recA^–^-R1 | GGGCGATACTATGAGAAGACC | 652 bp |
| recA^–^-F2 | GGGTCTTCTCATAGTATCGCC | recA^–^-R2 | CTGACAGTGAACTGATGCAG | 622 bp |
| recA^–^-F3 | GCGTAGAATTTCAGCGCGTTA | recA^–^-R3 | CTGACAGTGAACTGATGCAG | 1252 bp |
| ΔlacIZYA-F1 | GACTGTAGCGGCTGATGTT | ΔlacIZYA-R1 | ACGACATTGGCGTAAGTGAA | None |
| ΔlacIZYA-F2 | CCTCTTCCCACAAGACAACA | ΔlacIZYA-R2 | AGGTGCACCACGTTGTTTTA | 1323 bp |
| ΔNo.5::7.9kb-F1 | CGGTCGATATGCGGATGTAT | ΔNo.5::7.9kb-R1 | TCTCCTGCTGCAGGATTTTG | None |
| ΔNo.5::7.9kb-F2 | GTCGGTGTGTGTACGGTATT | ΔNo.5::7.9kb-R2 | CCGGCAGCTTGATATGTTTC | 1410 bp |
| ΔNo.5::7.9kb-F3 | GGTTTCAGTGGCTTGGTTCT | ΔNo.5::7.9kb-R3 | ACATACGGGAACTGCTCTTT | 1422 bp |
| RNAP(T7)-F | CTATCCTCTGCCGCTATGAA | RNAP(T7)-R | CGTTGCACATACGGGTTATG | 4322 bp |

**Table S3. Reagents and media used in this study**

| Reagent | Formulation concentration | Working concentration |
| --- | --- | --- |
| Ampicillin | 100 g/L | 0.1 g/L |
| Kanamycin | 50 g/L | 0.05 g/L |
| Glucose | 500 g/L | 10 g/L |
| IPTG | 100 mM | 1 mM |
| L-arabinose | 1 M | 20 mM/5 mM |
| Sucrose | 500 g/L | 20 g/L |
| X-gal | 20 g/L | 0.1 g/L |
| Medium | Composition | |
| LB | 10 g/L tryptone, 5 g/L yeast extract, 10 g/L NaCl; Solid medium (20 g/L agar) | |
| SOC | 20 g/L tryptone, 5 g/L yeast extract, 0.5 g/L NaCl, 2.5 mM KCl, 10 mM MgCl_2_, 10 mM MgSO_4_, 20 mM glucose | |
| M9 | 6 g/L Na_2_HPO_4_, 3 g/L KH_2_PO_4_, 0.5 g/L NaCl, 1 g/L NH_4_Cl, 1 mM MgSO_4_, 0.1 mM CaCl_2_, 10 mg/L VB_1_, 40 g/L glucose, 4 g/L yeast extract | |

**Table S4. Plasmids involved in this study**

| Plasmid | Description | Source/reference |
| --- | --- | --- |
| pUC19 | Used to measure transformation efficiency | [3] |
| p15A-P_araB_-Cas9-P_T5_-Redγβα | Plasmid#1 used in all genomic editing experiments | This study |
| pSC101-P_araB_-sgRNA-Donor-T1 | Parental plasmid for constructing plasmid#2 | This study |
| pSC101-P_araB_-sgRNA-Donor-T2 | Parental plasmid for constructing plasmid#2 | This study |
| pSC101-P_araB_-sgRNA-Donor-T3 | Parental plasmid for constructing plasmid#2 | This study |
| pSC101-P_araB_-sgRNA-Donor (*lacZ*::3kb) | Plasmid#2 used to insert 3-kb fragment into the *lacZ* gene | This study |
| pSC101-P_araB_-sgRNA-Donor (*lacZ*::6kb) | Plasmid#2 used to insert 6-kb fragment into the *lacZ* gene | This study |
| pSC101-P_araB_-sgRNA-Donor (*lacZ*::9kb) | Plasmid#2 used to insert 9-kb fragment into the *lacZ* gene | This study |
| pSC101-P_araB_-sgRNA-Donor (*lacZ*::12kb) | Plasmid#2 used to insert 12-kb fragment into the *lacZ* gene | This study |
| pSC101-P_araB_-sgRNA-Donor (Δ9.1kb) | Plasmid#2 used to delete the 9.1-kb fragment (565,156–574,260) | This study |
| pSC101-P_araB_-sgRNA-Donor (Δ21.5kb) | Plasmid#2 used to delete the 21.5-kb fragment (565,156–586,650) | This study |
| pSC101-P_araB_-sgRNA-Donor (Δ30.6kb) | Plasmid#2 used to delete the 30.6-kb fragment (565,156–595,752) | This study |
| pSC101-P_araB_-sgRNA-Donor (Δ39.4kb) | Plasmid#2 used to delete the 39.4-kb fragment (565,156–604,564) | This study |
| pSC101-P_araB_-sgRNA-Donor (Δ59.8kb) | Plasmid#2 used to delete the 59.8-kb fragment (565,156–624,919) | This study |
| pSC101-P_araB_-sgRNA-Donor (Δ79.8kb) | Plasmid#2 used to delete the 79.8-kb fragment (565,156–644,949) | This study |
| pSC101-P_araB_-sgRNA-Donor (Δ99.9kb/ΔNo.3) | Plasmid#2 used to delete the 99.9-kb fragment (fragment No.3) (565,156–665,089) | This study |
| pSC101-P_araB_-sgRNA-Donor (ΔNo.1) | Plasmid#2 used to delete the fragment No.1 (240,056–426,771) | This study |
| pSC101-P_araB_-sgRNA-Donor (ΔNo.2) | Plasmid#2 used to delete the fragment No.2 (499,529–552,955) | This study |
| pSC101-P_araB_-sgRNA-Donor (ΔNo.4) | Plasmid#2 used to delete the fragment No.4 (990,473–1,127,061) | This study |
| pSC101-P_araB_-sgRNA-Donor (ΔNo.5) | Plasmid#2 used to delete the fragment No.5 (1,449,596–1,549,490) | This study |
| pSC101-P_araB_-sgRNA-Donor (ΔNo.6) | Plasmid#2 used to delete the fragment No.6 (1,549,491–1,647,484) | This study |
| pSC101-P_araB_-sgRNA-Donor (ΔNo.7) | Plasmid#2 used to delete the fragment No.7 (2,349,152–2,430,141) | This study |
| pSC101-P_araB_-sgRNA-Donor (ΔNo.8) | Plasmid#2 used to delete the fragment No.8 (2,442,420–2,517,306) | This study |
| pSC101-P_araB_-sgRNA-Donor (ΔNo.9) | Plasmid#2 used to delete the fragment No.9 (2,822,534–2,906,555) | This study |
| pSC101-P_araB_-sgRNA-Donor (ΔNo.10) | Plasmid#2 used to delete the fragment No.10 (3,610,719–3,689,415) | This study |
| pSC101-P_araB_-sgRNA-Donor (ΔNo.11) | Plasmid#2 used to delete the fragment No.11 (3,824,765–3,876,879) | This study |
| pSC101-P_araB_-sgRNA-Donor (ΔNo.12) | Plasmid#2 used to delete the fragment No.12 (4,198,958–4,251,002) | This study |
| pSC101-P_araB_-sgRNA-Donor (ΔaraBAD::Tet^r^) | Plasmid#2 used to displace *araBAD* with Tet^r^ | This study |
| pSC101-P_araB_-sgRNA-Donor (Δ*mcrCB*-*hsdSMR*-*mrr*) | Plasmid#2 used to delete *mcrCB*-*hsdSMR*-*mrr* | This study |
| pSC101-P_araB_-sgRNA-Donor (*mcrA*^–^) | Plasmid#2 used to inactivate *mcrA* | This study |
| pSC101-P_araB_-sgRNA-Donor (*endA*^–^) | Plasmid#2 used to inactivate *endA* | This study |
| pSC101-P_araB_-sgRNA-Donor (*recA*^–^) | Plasmid#2 used to inactivate *recA* | This study |
| pSC101-P_araB_-sgRNA-Donor (Δ*lacIZYA*) | Plasmid#2 used to delete *lacIZYA* | This study |
| pSC101-P_araB_-sgRNA-Donor (Δfragment No.5::P_T7_-*alsS*-*ilvC*-*ilvD*-*kivD*-*adhA*-T_T7_) | Plasmid#2 used to displace fragment No.5 with P_T7_-*alsS*-*ilvC*-*ilvD*-*kivD*-*adhA*-T_T7_ | This study |
| pSC101-P_araB_-sgRNA-Donor (P_T5_-RNAP (T7)) | Plasmid#2 used to insert T7 RNA polymerase-encoding gene | This study |
| pColE1-P_T5_-*alsS*-*ilvC*-*ilvD*-*kivD*-*adhA* | Used to overexpress AlsS, IlvC, IlvD, KivD and AdhA | This study |

**Table S5. CRISPR target sequences designed in this study**

| Target site | Nucleotide sequence (5’-3’) | Target site | Nucleotide sequence (5’-3’) |
| --- | --- | --- | --- |
| lacZ | CAGTATCCCCGTTTACAGGGCGG | No.6-L | GAAGTGGCTAAAGAGAACAACGG |
| 9.1kb-L | GGTGTTTGTACATACCAAGGCGG | No.6-R | CAGCGTGTCACATATGAAGGTGG |
| 9.1kb-R | GAAAAGGTTACAGAAGGCTGAGG | No.7-L | TGGTGACTGAAGATATGGCGGGG |
| 21.5kb-L | GGTGTTTGTACATACCAAGGCGG | No.7-R | ATCTATGGTGATGATGCCGGCGG |
| 21.5kb-R | GCTATGACAAGCGATAACCGGGG | No.8-L | ATCCGGCGTCCTGCAAACCGCGG |
| 30.6kb-L | GGTGTTTGTACATACCAAGGCGG | No.8-R | CATGTTGTAAATCAGCACAGCGG |
| 30.6kb-R | GCAGATAACTATCAGAACGCGGG | No.9-L | AGAACTGCGACGTAAACTCGCGG |
| 39.4kb-L | GGTGTTTGTACATACCAAGGCGG | No.9-R | GGGGGCGGTTCTTCTGGAGGTGG |
| 39.4kb-R | AATAATATGCACCACGACGGCGG | No.10-L | TCCGCAGCAAAATCAGGGGGCGG |
| 59.8kb-L | GGTGTTTGTACATACCAAGGCGG | No.10-R | GCCACATCCACTTTTTCCGGCGG |
| 59.8kb-R | TGAGGAAGTACAGCAGACCATGG | No.11-L | CTGGCCTTTGAGATGAGCGACGG |
| 79.8kb-L | GGTGTTTGTACATACCAAGGCGG | No.11-R | GGCAATCGCATTTTTAACGGCGG |
| 79.8kb-R | TAAGGCGTTGGCAGAAGAGAAGG | No.12-L | TACTCGACCAGCTCAAGCGAGGG |
| 99.9kb-L | GGTGTTTGTACATACCAAGGCGG | No.12-R | TGATGGATAAAGATGCGCAGGGG |
| 99.9kb-R | ATAGGGCCGTTACATACCGCAGG | araBAD-L | GACCCGCCAGCTACAACCCGTGG |
| No.1-L | CTACCGGAGATGTTTACCAGCGG | araBAD-R | CAGAATCACTGCCAAAATCGAGG |
| No.1-R | TTTGATGGCTACGCTGTCGGCGG | mcrCB-hsdSMR-mrr-L | AAAATGGCGAAAACATAGGGGGG |
| No.2-L | TTATTTCCACCAGCAGGCGGAGG | mcrCB-hsdSMR-mrr-R | GCTGGGCAAAAAGCGAAACGTGG |
| No.2-R | GAAGCTGAACAGTTTATCGGGGG | mcrA | TGAGTGTTCGATAGGTGAAGAGG |
| No.4-L | AATGCCATCGACCAGAAAGGTGG | endA | TTATTTGTCTATTGCTGCGGTGG |
| No.4-R | CAGTACGTTACTCAATGCGGGGG | recA | CGAAAACAAACAGAAAGCGTTGG |
| No.5-L | GTAAAACCGAATACTGCCGGTGG | LacIZYA-L | ATCCGGCATGAACAAAGCGCAGG |
| No.5-R | GAAGTGGCTAAAGAGAACAACGG | LacIZYA-R | TTGTTCACTGCGCAGCGGTACGG |

**Table S6. Summary of optimized conditions**

| **Terms** | **Units** | **Tested conditions** | **Optimal condition** |
| --- | --- | --- | --- |
| Inducible promoter (Cas9, sgRNA) | － | P_araB_, P_T5_, P_L_lacO_1_ | P_araB_ |
| Inducible promoter 2 (λ-Red) | － | P_araB_, P_T5_, P_L_lacO_1_ | P_T5_ |
| Medium | － | SOB, SOC, LB, TB | LB |
| Time 1 | hour | 0.5, 1, 1.5, 2, 2.5, 3 | 2 |
| Time 2 | hour | 0.5, 1, 1.5, 2, 2.5, 3 | 1 |
| Time 3 | hour | 1, 2, 3, 4, 5 ,6 | 3 |
| L-arabinose (liquid) | mM | 1, 5, 10, 15, 20, 25, 30 | 20 |
| L-arabinose (plate) | mM | 1, 5, 10, 15, 20, 25, 30 | 5 |
| Temperature | ℃ | 25, 28, 30, 32, 35, 37 | 30 |

**Note S1. Complete sequence of plasmid p15A-P_araB_-Cas9-P_T5_-Redγβα in Genbank format**

LOCUS	p15A-P_araB_-Cas9-P_T5_-Redγβα	12867 bp	DNA	circular SYN 10-JAN-2020

DEFINITION p15A-P_araB_-Cas9-P_T5_-Redγβα

ACCESSION p15A-P_araB_-Cas9-P_T5_-Redγβα

KEYWORDS

SOURCE

ORGANISM

FEATURES	Location/Qualifiers

CDS	6..791

/label="Beta"

/note="single-stranded DNA binding recombinase in the λ Red system"

CDS	788..1468

/label="Exo"

/note="5' to 3' double-stranded DNA exonuclease in the λ Red system"

terminator	1469..1713

/label="lambda tL3 terminator"

/note="transcription terminator tL3 from phage lambda"

promoter	1745..1849

/label="AmpR promoter"

CDS	1850..2644

/label="KanR"

/note="aminoglycoside phosphotransferase from Tn5"

CDS	2661..3743

/label="lacI"

/note="lac repressor"

promoter	3770..4215

/label="sacB promoter"

CDS	4216..5637

/label="sacB"

/note="secreted levansucrase that renders bacterial growth sensitive to sucrose"

CDS	complement(5682..9788)

/label="Cas9"

/note="generates RNA-guided double strand breaks in DNA"

promoter	complement(9832..9997)

/label="araB promoter"

/note="promoter of the L-arabinose operon of *E. coli*"

CDS	10143..11021

/label="araC"

/note="L-arabinose regulatory protein"

terminator	11065..11111

/label="rrnB T1 terminator"

rep_origin	complement(11443..11988)

/label="p15A ori"

/note="replication origin"

promoter	12318..12367

/label="T5 promoter"

/note="bacteriophage T5 promoter for *E. coli* RNA polymerase, with embedded lac operator"

CDS	12451..12867

/label="Gam"

/note="inhibitor of the host RecBCD nuclease in the λ Red system"

ORIGIN

1 AACGAATGAG TACTGCACTC GCAACGCTGG CTGGGAAGCT GGCTGAACGT GTCGGCATGG

61 ATTCTGTCGA CCCACAGGAA CTGATCACCA CTCTTCGCCA GACGGCATTT AAAGGTGATG

121 CCAGCGATGC GCAGTTCATC GCATTACTGA TCGTTGCCAA CCAGTACGGC CTTAATCCGT

181 GGACGAAAGA AATTTACGCC TTTCCTGATA AGCAGAATGG CATCGTTCCG GTGGTGGGCG

241 TTGATGGCTG GTCCCGCATC ATCAATGAAA ACCAGCAGTT TGATGGCATG GACTTTGAGC

301 AGGACAATGA ATCCTGTACA TGCCGGATTT ACCGCAAGGA CCGTAATCAT CCGATCTGCG

361 TTACCGAATG GATGGATGAA TGCCGCCGCG AACCATTCAA AACTCGCGAA GGCAGAGAAA

421 TCACGGGGCC GTGGCAGTCG CATCCCAAAC GGATGTTACG TCATAAAGCC ATGATTCAGT

481 GTGCCCGTCT GGCCTTCGGA TTTGCTGGTA TCTATGACAA GGATGAAGCC GAGCGCATTG

541 TCGAAAATAC TGCATACACT GCAGAACGTC AGCCGGAACG CGACATCACT CCGGTTAACG

601 ATGAAACCAT GCAGGAGATT AACACTCTGC TGATCGCCCT GGATAAAACA TGGGATGACG

661 ACTTATTGCC GCTCTGTTCC CAGATATTTC GCCGCGACAT TCGTGCATCG TCAGAACTGA

721 CACAGGCCGA AGCAGTAAAA GCTCTTGGAT TCCTGAAACA GAAAGCCGCA GAGCAGAAGG

781 TGGCAGCATG ACACCGGACA TTATCCTGCA GCGTACCGGG ATCGATGTGA GAGCTGTCGA

841 ACAGGGGGAT GATGCGTGGC ACAAATTACG GCTCGGCGTC ATCACCGCTT CAGAAGTTCA

901 CAACGTGATA GCAAAACCCC GCTCCGGAAA GAAGTGGCCT GACATGAAAA TGTCCTACTT

961 CCACACCCTG CTTGCTGAGG TTTGCACCGG TGTGGCTCCG GAAGTTAACG CTAAAGCACT

1021 GGCCTGGGGA AAACAGTACG AGAACGACGC CAGAACCCTG TTTGAATTCA CTTCCGGCGT

1081 GAATGTTACT GAATCCCCGA TCATCTATCG CGACGAAAGT ATGCGTACCG CCTGCTCTCC

1141 CGATGGTTTA TGCAGTGACG GCAACGGCCT TGAACTGAAA TGCCCGTTTA CCTCCCGGGA

1201 TTTCATGAAG TTCCGGCTCG GTGGTTTCGA GGCCATAAAG TCAGCTTACA TGGCCCAGGT

1261 GCAGTACAGC ATGTGGGTGA CGCGAAAAAA TGCCTGGTAC TTTGCCAACT ATGACCCGCG

1321 TATGAAGCGT GAAGGCCTGC ATTATGTCGT GATTGAGCGG GATGAAAAGT ACATGGCGAG

1381 TTTTGACGAG ATCGTGCCGG AGTTCATCGA AAAAATGGAC GAGGCACTGG CTGAAATTGG

1441 TTTTGTATTT GGGGAGCAAT GGCGATGACG CATCCTCACG ATAATATCCG GGTAGGCGCA

1501 ATCACTTTCG TCTACTCCGT TACAAAGCGA GGCTGGGTAT TTCCCGGCCT TTCTGTTATC

1561 CGAAATCCAC TGAAAGCACA GCGGCTGGCT GAGGAGATAA ATAATAAACG AGGGGCTGTA

1621 TGCACAAAGC ATCTTCTGTT GAGTTAAGAA CGAGTATCGA GATGGCACAT AGCCTTGCTC

1681 AAATTGGAAT CAGGTTTGTG CCAATACCAG TAGAAACAGA CGAAGAATCC ATGGGTATGG

1741 ACAGCGCGGA ACCCCTATTT GTTTATTTTT CTAAATACAT TCAAATATGT ATCCGCTCAT

1801 GAGACAATAA CCCTGATAAA TGCTTCAATA ATATTGAAAA AGGAAGAGTA TGATTGAACA

1861 AGATGGATTG CACGCAGGTT CTCCGGCCGC TTGGGTGGAG AGGCTATTCG GCTATGACTG

1921 GGCACAACAG ACAATCGGCT GCTCTGATGC CGCCGTGTTC CGGCTGTCAG CGCAGGGGCG

1981 CCCGGTTCTT TTTGTCAAGA CCGACCTGTC CGGTGCCCTG AATGAACTGC AGGACGAGGC

2041 AGCGCGGCTA TCGTGGCTGG CCACGACGGG CGTTCCTTGC GCAGCTGTGC TCGACGTTGT

2101 CACTGAAGCG GGAAGGGACT GGCTGCTATT GGGCGAAGTG CCGGGGCAGG ATCTCCTGTC

2161 ATCTCACCTT GCTCCTGCCG AGAAAGTATC CATCATGGCT GATGCAATGC GGCGGCTGCA

2221 TACGCTTGAT CCGGCTACCT GCCCATTCGA CCACCAAGCG AAACATCGCA TCGAGCGAGC

2281 ACGTACTCGG ATGGAAGCCG GTCTTGTCGA TCAGGATGAT CTGGACGAAG AGCATCAGGG

2341 GCTCGCGCCA GCCGAACTGT TCGCCAGGCT CAAGGCGCGC ATGCCCGACG GCGAGGATCT

2401 CGTCGTGACC CATGGCGATG CCTGCTTGCC GAATATCATG GTGGAAAATG GCCGCTTTTC

2461 TGGATTCATC GACTGTGGCC GGCTGGGTGT GGCGGACCGC TATCAGGACA TAGCGTTGGC

2521 TACCCGTGAT ATTGCTGAAG AGCTTGGCGG CGAATGGGCT GACCGCTTCC TCGTGCTTTA

2581 CGGTATCGCC GCTCCCGATT CGCAGCGCAT CGCCTTCTAT CGCCTTCTTG ACGAGTTCTT

2641 CTGAAAGGAG GTTATAAAAA ATGAAACCAG TAACGTTATA CGATGTCGCA GAGTATGCCG

2701 GTGTCTCTTA TCAGACCGTT TCCCGCGTGG TGAACCAGGC CAGCCACGTT TCTGCGAAAA

2761 CGCGGGAAAA AGTGGAAGCG GCGATGGCGG AGCTGAATTA CATTCCCAAC CGCGTGGCAC

2821 AACAACTGGC GGGCAAACAG TCGTTGCTGA TTGGCGTTGC CACCTCCAGT CTGGCCCTGC

2881 ACGCGCCGTC GCAAATTGTC GCGGCGATTA AATCTCGCGC CGATCAACTG GGTGCCAGCG

2941 TGGTGGTGTC GATGGTAGAA CGAAGCGGCG TCGAAGCCTG TAAAGCGGCG GTGCACAATC

3001 TTCTCGCGCA ACGCGTCAGT GGGCTGATCA TTAACTATCC GCTGGATGAC CAGGATGCCA

3061 TTGCTGTGGA AGCTGCCTGC ACTAATGTTC CGGCGTTATT TCTTGATGTC TCTGACCAGA

3121 CACCCATCAA CAGTATTATT TTCTCCCATG AAGACGGTAC GCGACTGGGC GTGGAGCATC

3181 TGGTCGCATT GGGTCACCAG CAAATCGCGC TGTTAGCGGG CCCATTAAGT TCTGTCTCGG

3241 CGCGTCTGCG TCTGGCTGGC TGGCATAAAT ATCTCACTCG CAATCAAATT CAGCCGATAG

3301 CGGAACGGGA AGGCGACTGG AGTGCCATGT CCGGTTTTCA ACAAACCATG CAAATGCTGA

3361 ATGAGGGCAT CGTTCCCACT GCGATGCTGG TTGCCAACGA TCAGATGGCG CTGGGCGCAA

3421 TGCGCGCCAT TACCGAGTCC GGGCTGCGCG TTGGTGCGGA TATCTCGGTA GTGGGATACG

3481 ACGATACCGA AGACAGCTCA TGTTATATCC CGCCGTTAAC CACCATCAAA CAGGATTTTC

3541 GCCTGCTGGG GCAAACCAGC GTGGACCGCT TGCTGCAACT CTCTCAGGGC CAGGCGGTGA

3601 AGGGCAATCA GCTGTTGCCC GTCTCACTGG TGAAAAGAAA AACCACCCTG GCGCCCAATA

3661 CGCAAACCGC CTCTCCCCGC GCGTTGGCCG ATTCATTAAT GCAGCTGGCA CGACAGGTTT

3721 CCCGACTGGA AAGCGGGCAG TGATAACTGT CAGACCAAGT TTACGAGCTC ACATATACCT

3781 GCCGTTCACT ATTATTTAGT GAAATGAGAT ATTATGATAT TTTCTGAATT GTGATTAAAA

3841 AGGCAACTTT ATGCCCATGC AACAGAAACT ATAAAAAATA CAGAGAATGA AAAGAAACAG

3901 ATAGATTTTT TAGTTCTTTA GGCCCGTAGT CTGCAAATCC TTTTATGATT TTCTATCAAA

3961 CAAAAGAGGA AAATAGACCA GTTGCAATCC AAACGAGAGT CTAATAGAAT GAGGTCGAAA

4021 AGTAAATCGC GCGGGTTTGT TACTGATAAA GCAGGCAAGA CCTAAAATGT GTAAAGGGCA

4081 AAGTGTATAC TTTGGCGTCA CCCCTTACAT ATTTTAGGTC TTTTTTTATT GTGCGTAACT

4141 AACTTGCCAT CTTCAAACAG GAGGGCTGGA AGAAGCAGAC CGCTAACACA GTACATAAAA

4201 AAGGAGACAT GAACGATGAA CATCAAAAAG TTTGCAAAAC AAGCAACAGT ATTAACCTTT

4261 ACTACCGCAC TGCTGGCAGG AGGCGCAACT CAAGCGTTTG CGAAAGAAAC GAACCAAAAG

4321 CCATATAAGG AAACATACGG CATTTCCCAT ATTACACGCC ATGATATGCT GCAAATCCCT

4381 GAACAGCAAA AAAATGAAAA ATATCAAGTT CCTGAATTCG ATTCGTCCAC AATTAAAAAT

4441 ATCTCTTCTG CAAAAGGCCT GGACGTTTGG GACAGCTGGC CATTACAAAA CGCTGACGGC

4501 ACTGTCGCAA ACTATCACGG CTACCACATC GTCTTTGCAT TAGCCGGAGA TCCTAAAAAT

4561 GCGGATGACA CATCGATTTA CATGTTCTAT CAAAAAGTCG GCGAAACTTC TATTGACAGC

4621 TGGAAAAACG CTGGCCGCGT CTTTAAAGAC AGCGACAAAT TCGATGCAAA TGATTCTATC

4681 CTAAAAGACC AAACACAAGA ATGGTCAGGT TCAGCCACAT TTACATCTGA CGGAAAAATC

4741 CGTTTATTCT ACACTGATTT CTCCGGTAAA CATTACGGCA AACAAACACT GACAACTGCA

4801 CAAGTTAACG TATCAGCATC AGACAGCTCT TTGAACATCA ACGGTGTAGA GGATTATAAA

4861 TCAATCTTTG ACGGTGACGG AAAAACGTAT CAAAATGTAC AGCAGTTCAT CGATGAAGGC

4921 AACTACAGCT CAGGCGACAA CCATACGCTG AGAGATCCTC ACTACGTAGA AGATAAAGGC

4981 CACAAATACT TAGTATTTGA AGCAAACACT GGAACTGAAG ATGGCTACCA AGGCGAAGAA

5041 TCTTTATTTA ACAAAGCATA CTATGGCAAA AGCACATCAT TCTTCCGTCA AGAAAGTCAA

5101 AAACTTCTGC AAAGCGATAA AAAACGCACG GCTGAGTTAG CAAACGGCGC TCTCGGTATG

5161 ATTGAGCTAA ACGATGATTA CACACTGAAA AAAGTGATGA AACCGCTGAT TGCATCTAAC

5221 ACAGTAACAG ATGAAATTGA ACGCGCGAAC GTCTTTAAAA TGAACGGCAA ATGGTACCTG

5281 TTCACTGACT CCCGCGGATC AAAAATGACG ATTGACGGCA TTACGTCTAA CGATATTTAC

5341 ATGCTTGGTT ATGTTTCTAA TTCTTTAACT GGCCCATACA AGCCGCTGAA CAAAACTGGC

5401 CTTGTGTTAA AAATGGATCT TGATCCTAAC GATGTAACCT TTACTTACTC ACACTTCGCT

5461 GTACCTCAAG CGAAAGGAAA CAATGTCGTG ATTACAAGCT ATATGACAAA CAGAGGATTC

5521 TACGCAGACA AACAATCAAC GTTTGCGCCA AGCTTCCTGC TGAACATCAA AGGCAAGAAA

5581 ACATCTGTTG TCAAAGACAG CATCCTTGAA CAAGGACAAT TAACAGTTAA CAAATAAAAA

5641 CGCAAAAGAA AATGCCGATA TTGACTACCG GAAGCAGTGC TTCAGTCACC TCCTAGCTGA

5701 CTCAAATCAA TGCGTGTTTC ATAAAGACCA GTGATGGATT GATGGATAAG AGTGGCATCT

5761 AAAACTTCTT TTGTAGACGT ATATCGTTTA CGATCAATTG TTGTATCAAA ATATTTAAAA

5821 GCAGCGGGAG CTCCAAGATT CGTCAACGTA AATAAATGAA TAATATTTTC TGCTTGTTCA

5881 CGTATTGGTT TGTCTCTATG TTTGTTATAT GCACTAAGAA CTTTATCTAA ATTGGCATCT

5941 GCTAAAATAA CACGCTTAGA AAATTCACTG ATTTGCTCAA TAATCTCATC TAAATAATGC

6001 TTATGCTGCT CCACAAACAA TTGTTTTTGT TCGTTATCTT CTGGACTACC CTTCAACTTT

6061 TCATAATGAC TAGCTAAATA TAAAAAATTC ACATATTTGC TTGGCAGAGC CAGCTCATTT

6121 CCTTTTTGTA ATTCTCCGGC ACTAGCCAGC ATCCGTTTAC GACCGTTTTC TAACTCAAAA

6181 AGACTATATT TAGGTAGTTT AATGATTAAG TCTTTTTTAA CTTCCTTATA TCCTTTAGCT

6241 TCTAAAAAGT CAATCGGATT TTTTTCAAAG GAACTTCTTT CCATAATTGT GATCCCTAGT

6301 AACTCTTTAA CGGATTTTAA CTTCTTCGAT TTCCCTTTTT CCACCTTAGC AACCACTAGG

6361 ACTGAATAAG CTACCGTTGG ACTATCAAAA CCACCATATT TTTTTGGATC CCAGTCTTTT

6421 TTACGAGCAA TAAGCTTGTC CGAATTTCTT TTTGGTAAAA TTGACTCCTT GGAGAATCCG

6481 CCTGTCTGTA CTTCTGTTTT CTTGACAATA TTGACTTGGG GCATGGACAA TACTTTGCGC

6541 ACTGTGGCAA AATCTCGCCC TTTATCCCAG ACAATTTCTC CAGTTTCCCC ATTAGTTTCG

6601 ATTAGAGGGC GTTTGCGAAT CTCTCCATTT GCAAGTGTAA TTTCTGTTTT GAAGAAGTTC

6661 ATGATATTAG AGTAAAAGAA ATATTTTGCG GTTGCTTTGC CTATTTCTTG CTCAGACTTA

6721 GCAATCATTT TACGAACATC ATAAACTTTA TAATCACCAT AGACAAACTC CGATTCAAGT

6781 TTTGGATATT TCTTAATCAA AGCAGTTCCA ACGACGGCAT TTAGATACGC ATCATGGGCA

6841 TGATGGTAAT TGTTAATCTC ACGTACTTTA TAGAATTGGA AATCTTTTCG GAAGTCAGAA

6901 ACTAATTTAG ATTTTAAGGT AATCACTTTA ACCTCTCGAA TAAGTTTATC ATTTTCATCG

6961 TATTTAGTAT TCATGCGACT ATCCAAAATT TGTGCCACAT GCTTAGTGAT TTGGCGAGTT

7021 TCAACCAATT GGCGTTTGAT AAAACCAGCT TTATCAAGTT CACTCAAACC TCCACGTTCA

7081 GCTTTCGTTA AATTATCAAA CTTACGTTGA GTGATTAACT TGGCGTTTAG AAGTTGTCTC

7141 CAATAGTTTT TCATCTTTTT GACTACTTCT TCACTTGGAA CGTTATCCGA TTTACCACGA

7201 TTTTTATCAG AACGCGTTAA GACCTTATTG TCTATTGAAT CGTCTTTAAG GAAACTTTGT

7261 GGAACAATGT GATCGACATC ATAATCACTT AAACGATTAA TATCTAATTC TTGGTCCACA

7321 TACATGTCTC TTCCATTTTG GAGATAATAG AGATAGAGCT TTTCATTTTG CAATTGAGTA

7381 TTTTCAACAG GATGCTCTTT AAGAATCTGA CTTCCTAATT CTTTGATACC TTCTTCGATT

7441 CGTTTCATAC GCTCTCGCGA ATTTTTCTGG CCCTTTTGAG TTGTCTGATT TTCACGTGCC

7501 ATTTCAATAA CGATATTTTC TGGCTTATGC CGCCCCATTA CTTTGACCAA TTCATCAACA

7561 ACTTTTACAG TCTGTAAAAT ACCTTTTTTA ATAGCAGGGC TACCAGCTAA ATTTGCAATA

7621 TGTTCATGTA AACTATCGCC TTGTCCAGAC ACTTGTGCTT TTTGAATGTC TTCTTTAAAT

7681 GTCAAACTAT CATCATGGAT CAGCTGCATA AAATTGCGAT TGGCAAAACC ATCTGATTTC

7741 AAAAAATCTA ATATTGTTTT GCCAGATTGC TTATCCCTAA TACCATTAAT CAATTTTCGA

7801 GACAAACGTC CCCAACCAGT ATAACGGCGA CGTTTAAGCT GTTTCATCAC CTTATCATCA

7861 AAGAGGTGAG CATATGTTTT AAGTCTTTCC TCAATCATCT CCCTATCTTC AAATAAGGTC

7921 AATGTTAAAA CAATATCCTC TAAGATATCT TCATTTTCTT CATTATCCAA AAAATCTTTA

7981 TCTTTAATAA TTTTTAGCAA ATCATGGTAG GTACCTAATG AAGCATTAAA TCTATCTTCA

8041 ACTCCTGAAA TTTCAACACT ATCAAAACAT TCTATTTTTT TGAAATAATC TTCTTTTAAT

8101 TGCTTAACGG TTACTTTTCG ATTTGTTTTG AAGAGTAAAT CAACAATGGC TTTCTTCTGT

8161 TCACCTGAAA GAAATGCTGG TTTTCGCATT CCTTCAGTAA CATATTTGAC CTTTGTCAAT

8221 TCGTTATAAA CCGTAAAATA CTCATAAAGC AAACTATGTT TTGGTAGTAC TTTTTCATTT

8281 GGAAGATTTT TATCAAAGTT TGTCATGCGT TCAATAAATG ATTGAGCTGA AGCACCTTTA

8341 TCGACAACTT CTTCAAAATT CCATGGGGTA ATTGTTTCTT CAGACTTCCG AGTCATCCAT

8401 GCAAAACGAC TATTGCCACG CGCCAATGGA CCAACATAAT AAGGAATTCG AAAAGTCAAG

8461 ATTTTTTCAA TCTTCTCACG ATTGTCTTTT AAAAATGGAT AAAAGTCTTC TTGTCTTCTC

8521 AAAATAGCAT GCAGCTCACC CAAGTGAATT TGATGGGGAA TAGAGCCGTT GTCAAAGGTC

8581 CGTTGCTTGC GCAGCAAATC TTCACGATTT AGTTTCACCA ATAATTCCTC AGTACCATCC

8641 ATTTTTTCTA AAATTGGTTT GATAAATTTA TAAAATTCTT CTTGGCTAGC TCCCCCATCA

8701 ATATAACCTG CATATCCGTT TTTTGATTGA TCAAAAAAGA TTTCTTTATA CTTTTCTGGA

8761 AGTTGTTGTC GAACTAAAGC TTTTAAAAGA GTCAAGTCTT GATGATGTTC ATCGTAGCGT

8821 TTAATCATTG AAGCTGATAG GGGAGCCTTA GTTATTTCAG TATTTACTCT TAGGATATCT

8881 GAAAGTAAAA TAGCATCTGA TAAATTCTTA GCTGCCAAAA ACAAATCAGC ATATTGATCT

8941 CCAATTTGCG CCAATAAATT ATCTAAATCA TCATCGTAAG TATCTTTTGA AAGCTGTAAT

9001 TTAGCATCTT CTGCCAAATC AAAATTTGAT TTAAAATTAG GGGTCAAACC CAATGACAAA

9061 GCAATGAGAT TCCCAAATAA GCCATTTTTC TTCTCACCGG GGAGCTGAGC AATGAGATTT

9121 TCTAATCGTC TTGATTTACT CAATCGTGCA GAAAGAATCG CTTTAGCATC TACTCCACTT

9181 GCGTTAATAG GGTTTTCTTC AAATAATTGA TTGTAGGTTT GTACCAACTG GATAAATAGT

9241 TTGTCCACAT CACTATTATC AGGATTTAAA TCTCCCTCAA TCAAAAAATG ACCACGAAAC

9301 TTAATCATAT GCGCTAAGGC CAAATAGATT AAGCGCAAAT CCGCTTTATC AGTAGAATCT

9361 ACCAATTTTT TTCGCAGATG ATAGATAGTT GGATATTTCT CATGATAAGC AACTTCATCT

9421 ACTATATTTC CAAAAATAGG ATGACGTTCA TGCTTCTTGT CTTCTTCCAC CAAAAAAGAC

9481 TCTTCAAGTC GATGAAAGAA ACTATCATCT ACTTTCGCCA TCTCATTTGA AAAAATCTCC

9541 TGTAGATAAC AAATACGATT CTTCCGACGT GTATACCTTC TACGAGCTGT CCGTTTGAGA

9601 CGAGTCGCTT CCGCTGTCTC TCCACTGTCA AATAAAAGAG CCCCTATAAG ATTTTTTTTG

9661 ATACTGTGGC GGTCTGTATT TCCCAGAACC TTGAACTTTT TAGACGGAAC CTTATATTCA

9721 TCAGTGATCA CCGCCCATCC GACGCTATTT GTGCCGATAT CTAAGCCTAT TGAGTATTTC

9781 TTATCCATTT TTTATAACCT CCTTAGAGCT CGAATTCCCA AAAAAACGGG TATGGAGAAA

9841 CAGTAGAGAG TTGCGATAAA AAGCGTCAGG TAGGATCCGC TAATCTTATG GATAAAAATG

9901 CTATGGCATA GCAAAGTGTG ACGCCGTGCA AATAATCAAT GTGGACTTTT CTGCCGTGAT

9961 TATAGACACT TTTGTTACGC GTTTTTGTCA TGGCTTTGGT CCCGCTTTGT TACAGAATGC

10021 TTTTAATAAG CGGGGTTACC GGTTTGGTTA GCGAGAAGAG CCAGTAAAAG ACGCAGTGAC

10081 GGCAATGTCT GATGCAATAT GGACAATTGG TTTCTTCTCT GAATGGCGGG AGTATGAAAA

10141 GTATGGCTGA AGCGCAAAAT GATCCCCTGC TGCCGGGATA CTCGTTTAAT GCCCATCTGG

10201 TGGCGGGTTT AACGCCGATT GAGGCCAACG GTTATCTCGA TTTTTTTATC GACCGACCGC

10261 TGGGAATGAA AGGTTATATT CTCAATCTCA CCATTCGCGG TCAGGGGGTG GTGAAAAATC

10321 AGGGACGAGA ATTTGTTTGC CGACCGGGTG ATATTTTGCT GTTCCCGCCA GGAGAGATTC

10381 ATCACTACGG TCGTCATCCG GAGGCTCGCG AATGGTATCA CCAGTGGGTT TACTTTCGTC

10441 CGCGCGCCTA CTGGCATGAA TGGCTTAACT GGCCGTCAAT ATTTGCCAAT ACGGGGTTCT

10501 TTCGCCCGGA TGAAGCGCAC CAGCCGCATT TCAGCGACCT GTTTGGGCAA ATCATTAACG

10561 CCGGGCAAGG GGAAGGGCGC TATTCGGAGC TGCTGGCGAT AAATCTGCTT GAGCAATTGT

10621 TACTGCGGCG CATGGAAGCG ATTAACGAGT CGCTCCATCC ACCGATGGAT AATCGGGTAC

10681 GCGAGGCTTG TCAGTACATC AGCGATCACC TGGCAGACAG CAATTTTGAT ATCGCCAGCG

10741 TCGCACAGCA TGTTTGCTTG TCGCCGTCGC GTCTGTCACA TCTTTTCCGC CAGCAGTTAG

10801 GGATTAGCGT CTTAAGCTGG CGCGAGGACC AACGTATCAG CCAGGCGAAG CTGCTTTTGA

10861 GCACCACCCG GATGCCTATC GCCACCGTCG GTCGCAATGT TGGTTTTGAC GATCAACTCT

10921 ATTTCTCGCG GGTATTTAAA AAATGCACCG GGGCCAGCCC GAGCGAGTTC CGTGCCGGTT

10981 GTGAAGAAAA AGTGAATGAT GTAGCCGTCA AGTTGTCATA ATAAATCGAT GCAGGTGGCA

11041 CTTTTCGGGG AAATGTGGAG GCATCAAATA AAACGAAAGG CTCAGTCGAA AGACTGGGCC

11101 TTTCGTTTTA TCTGTTGTTT GTCGGTGAAC GCTCTCCTGA GTAGGACAAA TCCGCCGCCC

11161 TAGACCTAGG GCGTTCGGCT GCGGCGAGCG GTATCAGCTC ACTCAAAGGC GGTAATACGG

11221 TTATCCACAG AATCAGGGGA TAACGCAGGA AAGAGCATGT GAGCAAAAGG CCAGCAAAAG

11281 GCCAGGAACC GTGGATATAT TCCGCTTCCT CGCTCACTGA CTCGCTACGC TCGGTCGTTC

11341 GACTGCGGCG AGCGGAAATG GCTTACGAAC GGGGCGGAGA TTTCCTGGAA GATGCCAGGA

11401 AGATACTTAA CAGGGAAGTG AGAGGGCCGC GGCAAAGCCG TTTTTCCATA GGCTCCGCCC

11461 CCCTGACAAG CATCACGAAA TCTGACGCTC AAATCAGTGG TGGCGAAACC CGACAGGACT

11521 ATAAAGATAC CAGGCGTTTC CCCCTGGCGG CTCCCTCGTG CGCTCTCCTG TTCCTGCCTT

11581 TCGGTTTACC GGTGTCATTC CGCTGTTATG GCCGCGTTTG TCTCATTCCA CGCCTGACAC

11641 TCAGTTCCGG GTAGGCAGTT CGCTCCAAGC TGGACTGTAT GCACGAACCC CCCGTTCAGT

11701 CCGACCGCTG CGCCTTATCC GGTAACTATC GTCTTGAGTC CAACCCGGAA AGACATGCAA

11761 AAGCACCACT GGCAGCAGCC ACTGGTAATT GATTTAGAGG AGTTAGTCTT GAAGTCATGC

11821 GCCGGTTAAG GCTAAACTGA AAGGACAAGT TTTGGTGACT GCGCTCCTCC AAGCCAGTTA

11881 CCTCGGTTCA AAGAGTTGGT AGCTCAGAGA ACCTTCGAAA AACCGCCCTG CAAGGCGGTT

11941 TTTTCGTTTT CAGAGCAAGA GATTACGCGC AGACCAAAAC GATCTCAAGA AGATCATCTT

12001 ATTAATAAGG ATCTCAAGAA GATCCTTTGA TCTTTTCTAC GGGGTCTGAC GCTCAGTGGA

12061 ACGAAAACTC ACGTTAAGGG ATTTTGGTCA TGACTAGTGC TTGGATTCTC ACCAATAAAA

12121 AACGCCCGGC GGCAACCGAT TTCAAGTTGA TAACGGACTA GCCTTATTTT AACTTGCTAT

12181 GCTGTTTTGA ATGGTTCCAA CAAGATTATT TTATAACTTT TATAACAAAT AATCAAGGAG

12241 AAATTCAAAG AAATTTATCA GCCGTGTCGC CCTTAATTGT GAGCGGATAA CAATTACGAG

12301 CTTCATGCAC AGTGAAATCA TGAAAAATTT ATTTGCTTTG TGAGCGGATA ACAATTATAA

12361 TATGTGGAAT TGTGAGCGCT CACAATTCCA CAACGGTTTC CCTCTAGAAA TAATTTTGTT

12421 TAACTTTTCG AGACCTTAGG AGGTAAACAT ATGGATATTA ATACTGAAAC TGAGATCAAG

12481 CAAAAGCATT CACTAACCCC CTTTCCTGTT TTCCTAATCA GCCCGGCATT TCGCGGGCGA

12541 TATTTTCACA GCTATTTCAG GAGTTCAGCC ATGAACGCTT ATTACATTCA GGATCGTCTT

12601 GAGGCTCAGA GCTGGGCGCG TCACTACCAG CAGCTCGCCC GTGAAGAGAA AGAGGCAGAA

12661 CTGGCAGACG ACATGGAAAA AGGCCTGCCC CAGCACCTGT TTGAATCGCT ATGCATCGAT

12721 CATTTGCAAC GCCACGGGGC CAGCAAAAAA TCCATTACCC GTGCGTTTGA TGACGATGTT

12781 GAGTTTCAGG AGCGCATGGC AGAACACATC CGGTACATGG TTGAAACCAT TGCTCACCAC

12841 CAGGTTGATA TTGATTCAGA GGTATAA

//

**Note S2. Complete sequence of plasmid pSC101-P_araB_-sgRNA-Donor-T1 in Genbank format**

LOCUS	pSC101-P_araB_-sgRNA-Donor-T1	3893 bp	DNA	circular SYN 10-JAN-2020

DEFINITION	pSC101-P_araB_-sgRNA-Donor-T1

ACCESSION	pSC101-P_araB_-sgRNA-Donor-T1

KEYWORDS

SOURCE

ORGANISM

FEATURES	Location/Qualifiers

CDS	complement(1..951)

/label="rep101"

/note="replication protein for the pSC101 origin"

rep_origin	999..1221

/label="pSC101 ori"

/note=" replication origin"

CDS	complement(1843..2703)

/label="AmpR"

/note="ampicillin resistance gene"

promoter	complement(2704..2808)

/label="AmpR promoter"

misc_feature	3230..3247

/label="I-SceI"

/note="recognition sequence of homing endonuclease I-SceI"

promoter	3393..3558

/label="araBAD promoter"

misc_RNA	3559..3634

/label="gRNA scaffold"

/note="guide RNA scaffold for the CRISPR/Cas9 system"

terminator	3695..3766

/label="rrnB T1 terminator"

ORIGIN

1 TCAGATCCTT CCGTATTTAG CCAGTATGTT CTCTAGTGTG GTTCGTTGTT TTTGCGTGAG

61 CCATGAGAAC GAACCATTGA GATCATACTT ACTTTGCATG TCACTCAAAA ATTTTGCCTC

121 AAAACTGGTG AGCTGAATTT TTGCAGTTAA AGCATCGTGT AGTGTTTTTC TTAGTCCGTT

181 ACGTAGGTAG GAATCTGATG TAATGGTTGT TGGTATTTTG TCACCATTCA TTTTTATCTG

241 GTTGTTCTCA AGTTCGGTTA CGAGATCCAT TTGTCTATCT AGTTCAACTT GGAAAATCAA

301 CGTATCAGTC GGGCGGCCTC GCTTATCAAC CACCAATTTC ATATTGCTGT AAGTGTTTAA

361 ATCTTTACTT ATTGGTTTCA AAACCCATTG GTTAAGCCTT TTAAACTCAT GGTAGTTATT

421 TTCAAGCATT AACATGAACT TAAATTCATC AAGGCTAATC TCTATATTTG CCTTGTGAGT

481 TTTCTTTTGT GTTAGTTCTT TTAATAACCA CTCATAAATC CTCATAGAGT ATTTGTTTTC

541 AAAAGACTTA ACATGTTCCA GATTATATTT TATGAATTTT TTTAACTGGA AAAGATAAGG

601 CAATATCTCT TCACTAAAAA CTAATTCTAA TTTTTCGCTT GAGAACTTGG CATAGTTTGT

661 CCACTGGAAA ATCTCAAAGC CTTTAACCAA AGGATTCCTG ATTTCCACAG TTCTCGTCAT

721 CAGCTCTCTG GTTGCTTTAG CTAATACACC ATAAGCATTT TCCCTACTGA TGTTCATCAT

781 CTGAGCGTAT TGGTTATAAG TGAACGATAC CGTCCGTTCT TTCCTTGTAG GGTTTTCAAT

841 CGTGGGGTTG AGTAGTGCCA CACAGCATAA AATTAGCTTG GTTTCATGCT CCGTTAAGTC

901 ATAGCGACTA ATCGCTAGTT CATTTGCTTT GAAAACAACT AATTCAGACA TACATCTCAA

961 TTGGTCTAGG TGATTTTAAT CACTATACCA ATTGAGATGG GCTAGTCAAT GATAATTACT

1021 AGTCCTTTTC CTTTGAGTTG TGGGTATCTG TAAATTCTGC TAGACCTTTG CTGGAAAACT

1081 TGTAAATTCT GCTAGACCCT CTGTAAATTC CGCTAGACCT TTGTGTGTTT TTTTTGTTTA

1141 TATTCAAGTG GTTATAATTT ATAGAATAAA GAAAGAATAA AAAAAGATAA AAAGAATAGA

1201 TCCCAGCCCT GTGTATAACT CACTACTTTA GTCAGTTCCG CAGTATTACA AAAGGATGTC

1261 GCAAACGCTG TTTGCTCCTC TACAAAACAG ACCTTAAAAC CCTAAAGGCT TAAGTAGCAC

1321 CCTCGCAAGC TCGGTTGCGG CCGCAATCGG GCAAATCGCT GAATATTCCT TTTGTCTCCG

1381 ACCATCAGGC ACCTGAGTCG CTGTCTTTTT CGTGACATTC AGTTCGCTGC GCTCACGGCT

1441 CTGGCAGTGA ATGGGGGTAA ATGGCACTAC AGGCGCCTTT TATGGATTCA TGCAAGGAAA

1501 CTACCCATAA TACAAGAAAA GCCCGTCACG GGCTTCTCAG GGCGTTTTAT GGCGGGTCTG

1561 CTATGTGGTG CTATCTGACT TTTTGCTGTT CAGCAGTTCC TGCCCTCTGA TTTTCCAGTC

1621 TGACCACTTC GGATTATCCC GTGACAGGTC ATTCAGACTG GCTAATGCAC CCAGTAAGGC

1681 AGCGGTATCA TCAACGGGGT CTGACGCTCA GTGGAACGAA AACTCACGTT AAGGGATTTT

1741 GGTCATGAGA TTATCAAAAA GGATCTTCAC CTAGATCCTT TTAAATTAAA AATGAAGTTT

1801 TAAATCAATC TAAAGTATAT ATGAGTAAAC TTGGTCTGAC AGTTACCAAT GCTTAATCAG

1861 TGAGGCACCT ATCTCAGCGA TCTGTCTATT TCGTTCATCC ATAGTTGCCT GACTCCCCGT

1921 CGTGTAGATA ACTACGATAC GGGAGGGCTT ACCATCTGGC CCCAGTGCTG CAATGATACC

1981 GCGAGACCCA CGCTCACCGG CTCCAGATTT ATCAGCAATA AACCAGCCAG CCGGAAGGGC

2041 CGAGCGCAGA AGTGGTCCTG CAACTTTATC CGCCTCCATC CAGTCTATTA ATTGTTGCCG

2101 GGAAGCTAGA GTAAGTAGTT CGCCAGTTAA TAGTTTGCGC AACGTTGTTG CCATTGCTAC

2161 AGGCATCGTG GTGTCACGCT CGTCGTTTGG TATGGCTTCA TTCAGCTCCG GTTCCCAACG

2221 ATCAAGGCGA GTTACATGAT CCCCCATGTT GTGCAAAAAA GCGGTTAGCT CCTTCGGTCC

2281 TCCGATCGTT GTCAGAAGTA AGTTGGCCGC AGTGTTATCA CTCATGGTTA TGGCAGCACT

2341 GCATAATTCT CTTACTGTCA TGCCATCCGT AAGATGCTTT TCTGTGACTG GTGAGTACTC

2401 AACCAAGTCA TTCTGAGAAT AGTGTATGCG GCGACCGAGT TGCTCTTGCC CGGCGTCAAT

2461 ACGGGATAAT ACCGCGCCAC ATAGCAGAAC TTTAAAAGTG CTCATCATTG GAAAACGTTC

2521 TTCGGGGCGA AAACTCTCAA GGATCTTACC GCTGTTGAGA TCCAGTTCGA TGTAACCCAC

2581 TCGTGCACCC AACTGATCTT CAGCATCTTT TACTTTCACC AGCGTTTCTG GGTGAGCAAA

2641 AACAGGAAGG CAAAATGCCG CAAAAAAGGG AATAAGGGCG ACACGGAAAT GTTGAATACT

2701 CATACTCTTC CTTTTTCAAT ATTATTGAAG CATTTATCAG GGTTATTGTC TCATGAGCGG

2761 ATACATATTT GAATGTATTT AGAAAAATAA ACAAATAGGG GTTCCGCGGC ACAGATGCGT

2821 AAGGAGAAAA TACCGCATCA GGCGCCATTC GCCATTCAGG CTGCGCAACT GTTGGGAAGG

2881 GCGATCGGTG CGGGCCTCTT CGCTATTACG CCAGCTGGCG AAAGGGGGAT GTGCTGCAAG

2941 GCGATTAAGT TGGGTAACGC CAGGGTTTTC CCAGTCACGA CGTTGTAAAA CGACGGCCAG

3001 TGCCAAGCTT GCATGCCTGC AGGTCGACTC TAGAGGATCC CCGGGTACCG AGCTCGAATT

3061 CGTAATCATG TCATAGCTGT TTCCTGTGTG AAATTGTTAT CCGCTCACAA TTCCACACAA

3121 CATACGAGCC GGAAGCATAA AGTGTAAAGC CTGGGGTGCC TAATGAGTGA GCTAACTCAC

3181 ATTAATTGCG TTGCGCTCAC TGCCCGCTTT CCAGTCGGGA AACCTGTCAT AGGGATAACA

3241 GGGTAATACT TTTCATACTC CCGCCATTCA GAGAAGAAAC CAATTGTCCA TATTGCATCA

3301 GACATTGCCG TCACTGCGTC TTTTACTGGC TCTTCTCGCT AACCAAACCG GTAACCCCGC

3361 TTATTAAAAG CATTCTGTAA CAAAGCGGGA CCAAAGCCAT GACAAAAACG CGTAACAAAA

3421 GTGTCTATAA TCACGGCAGA AAAGTCCACA TTGATTATTT GCACGGCGTC ACACTTTGCT

3481 ATGCCATAGC ATTTTTATCC ATAAGATTAG CGGATCCTAC CTGACGCTTT TTATCGCAAC

3541 TCTCTACTGT TTCTCCATGT TTTAGAGCTA GAAATAGCAA GTTAAAATAA GGCTAGTCCG

3601 TTATCAACTT GAAAAAGTGG CACCGAGTCG GTGCTTAGCA TCCAAACTCG AGTAAGGATC

3661 ATTAAGGATC CCATGGTACG CGTGCTAGAG GCATCAAATA AAACGAAAGG CTCAGTCGAA

3721 AGACTGGGCC TTTCGTTTTA TCTGTTGTTT GTCGGTGAAC GCTCTCCTGA GTAGGACAAA

3781 TCCGCCCCAT GGGTATGGAC AGTTTTCCCT TTGATATGTA ACGGTGAACA GTTGTTCTAC

3841 TTTTGTTTGT TAGTCTTGAT GCTTCACTGA TAGATACAAG AGCCATAAGA ACC

//

**Note S3. Complete sequence of plasmid pSC101-P_araB_-sgRNA-Donor-T2 in Genbank format**

LOCUS	pSC101-P_araB_-sgRNA-Donor-T2	2998 bp	DNA	circular SYN 10-JAN-2020

DEFINITION	pSC101-P_araB_-sgRNA-Donor-T2

ACCESSION	pSC101-P_araB_-sgRNA-Donor-T2

KEYWORDS

SOURCE

ORGANISM

FEATURES	Location/Qualifiers

promoter	96..200

/label="AmpR promoter"

CDS	201..1061

/label="AmpR"

/note="ampicillin resistance gene"

promoter	1339..1504

/label="araBAD promoter"

rep_origin	1539..2127

/label="colE1 ori"

/note="replication origin"

misc_RNA	2727..2802

/label="gRNA scaffold"

/note="guide RNA scaffold for the CRISPR/Cas9 system"

terminator	2863..2934

/label="rrnB T1 terminator"

ORIGIN

1 GACGAAAGGG CCTCGTGATA CGCCTATTTT TATAGGTTAA TGTCATGATA ATAATGGTTT

61 CTTAGACGTC AGGTGGCACT TTTCGGGGAA ATGTGCGCGG AACCCCTATT TGTTTATTTT

121 TCTAAATACA TTCAAATATG TATCCGCTCA TGAGACAATA ACCCTGATAA ATGCTTCAAT

181 AATATTGAAA AAGGAAGAGT ATGAGTATTC AACATTTCCG TGTCGCCCTT ATTCCCTTTT

241 TTGCGGCATT TTGCCTTCCT GTTTTTGCTC ACCCAGAAAC GCTGGTGAAA GTAAAAGATG

301 CTGAAGATCA GTTGGGTGCA CGAGTGGGTT ACATCGAACT GGATCTCAAC AGCGGTAAGA

361 TCCTTGAGAG TTTTCGCCCC GAAGAACGTT TTCCAATGAT GAGCACTTTT AAAGTTCTGC

421 TATGTGGCGC GGTATTATCC CGTATTGACG CCGGGCAAGA GCAACTCGGT CGCCGCATAC

481 ACTATTCTCA GAATGACTTG GTTGAGTACT CACCAGTCAC AGAAAAGCAT CTTACGGATG

541 GCATGACAGT AAGAGAATTA TGCAGTGCTG CCATAACCAT GAGTGATAAC ACTGCGGCCA

601 ACTTACTTCT GACAACGATC GGAGGACCGA AGGAGCTAAC CGCTTTTTTG CACAACATGG

661 GGGATCATGT AACTCGCCTT GATCGTTGGG AACCGGAGCT GAATGAAGCC ATACCAAACG

721 ACGAGCGTGA CACCACGATG CCTGTAGCAA TGGCAACAAC GTTGCGCAAA CTATTAACTG

781 GCGAACTACT TACTCTAGCT TCCCGGCAAC AATTAATAGA CTGGATGGAG GCGGATAAAG

841 TTGCAGGACC ACTTCTGCGC TCGGCCCTTC CGGCTGGCTG GTTTATTGCT GATAAATCTG

901 GAGCCGGTGA GCGTGGGTCT CGCGGTATCA TTGCAGCACT GGGGCCAGAT GGTAAGCCCT

961 CCCGTATCGT AGTTATCTAC ACGACGGGGA GTCAGGCAAC TATGGATGAA CGAAATAGAC

1021 AGATCGCTGA GATAGGTGCC TCACTGATTA AGCATTGGTA ACTGTCAGAC CAAGTTTACT

1081 CATATATACT TTAGATTGAT TTAAAACTTC ATTTTTAATT TAAAAGGATC TAGGTGAAGA

1141 TCCTTTTTGA TAATCTCATG ACCAAAATCC CTTAACGTGA GTTTTCGTTC CACACTTTTC

1201 ATACTCCCGC CATTCAGAGA AGAAACCAAT TGTCCATATT GCATCAGACA TTGCCGTCAC

1261 TGCGTCTTTT ACTGGCTCTT CTCGCTAACC AAACCGGTAA CCCCGCTTAT TAAAAGCATT

1321 CTGTAACAAA GCGGGACCAA AGCCATGACA AAAACGCGTA ACAAAAGTGT CTATAATCAC

1381 GGCAGAAAAG TCCACATTGA TTATTTGCAC GGCGTCACAC TTTGCTATGC CATAGCATTT

1441 TTATCCATAA GATTAGCGGA TCCTACCTGA CGCTTTTTAT CGCAACTCTC TACTGTTTCT

1501 CCATCGTCAG ACCCCGTAGA AAAGATCAAA GGATCTTCTT GAGATCCTTT TTTTCTGCGC

1561 GTAATCTGCT GCTTGCAAAC AAAAAAACCA CCGCTACCAG CGGTGGTTTG TTTGCCGGAT

1621 CAAGAGCTAC CAACTCTTTT TCCGAAGGTA ACTGGCTTCA GCAGAGCGCA GATACCAAAT

1681 ACTGTCCTTC TAGTGTAGCC GTAGTTAGGC CACCACTTCA AGAACTCTGT AGCACCGCCT

1741 ACATACCTCG CTCTGCTAAT CCTGTTACCA GTGGCTGCTG CCAGTGGCGA TAAGTCGTGT

1801 CTTACCGGGT TGGACTCAAG ACGATAGTTA CCGGATAAGG CGCAGCGGTC GGGCTGAACG

1861 GGGGGTTCGT GCACACAGCC CAGCTTGGAG CGAACGACCT ACACCGAACT GAGATACCTA

1921 CAGCGTGAGC TATGAGAAAG CGCCACGCTT CCCGAAGGGA GAAAGGCGGA CAGGTATCCG

1981 GTAAGCGGCA GGGTCGGAAC AGGAGAGCGC ACGAGGGAGC TTCCAGGGGG AAACGCCTGG

2041 TATCTTTATA GTCCTGTCGG GTTTCGCCAC CTCTGACTTG AGCGTCGATT TTTGTGATGC

2101 TCGTCAGGGG GGCGGAGCCT ATGGAAAAAC GCCAGCAACG CGGCCTTTTT ACGGTTCCTG

2161 GCCTTTTGCT GGCCTTTTGC TCACATGTTC TTTCCTGCGT TATCCCCTGA TTCTGTGGAT

2221 AACCGTATTA CCGCCTTTGA GTGAGCTGAT ACCGCTGCAC AGATGCGTAA GGAGAAAATA

2281 CCGCATCAGG CGCCATTCGC CATTCAGGCT GCGCAACTGT TGGGAAGGGC GATCGGTGCG

2341 GGCCTCTTCG CTATTACGCC AGCTGGCGAA AGGGGGATGT GCTGCAAGGC GATTAAGTTG

2401 GGTAACGCCA GGGTTTTCCC AGTCACGACG TTGTAAAACG ACGGCCAGTG CCAAGCTTGC

2461 ATGCCTGCAG GTCGACTCTA GAGGATCCCC GGGTACCGAG CTCGAATTCG TAATCATGTC

2521 ATAGCTGTTT CCTGTGTGAA ATTGTTATCC GCTCACAATT CCACACAACA TACGAGCCGG

2581 AAGCATAAAG TGTAAAGCCT GGGGTGCCTA ATGAGTGAGC TAACTCACAT TAATTGCGTT

2641 GCGCTCACTG CCCGCTTTCC AGTCGGGAAA CCTGTCATAA CACCGTGCGT GTTGACTATT

2701 TTACCTCTGG CGGTGATAAT GGTTGCGTTT TAGAGCTAGA AATAGCAAGT TAAAATAAGG

2761 CTAGTCCGTT ATCAACTTGA AAAAGTGGCA CCGAGTCGGT GCTTAGCATC CAAACTCGAG

2821 TAAGGATCAT TAAGGATCCC ATGGTACGCG TGCTAGAGGC ATCAAATAAA ACGAAAGGCT

2881 CAGTCGAAAG ACTGGGCCTT TCGTTTTATC TGTTGTTTGT CGGTGAACGC TCTCCTGAGT

2941 AGGACAAATC CGCCCTGCAT GTGTCAGAGG TTTTCACCGT CATCACCGAA ACGCGCGA

//

**Note S3. Complete sequence of plasmid pSC101-P_araB_-sgRNA-Donor-T3 in Genbank format**

LOCUS	pSC101-P_araB_-sgRNA-Donor-T3	3942 bp	DNA	circular SYN 10-JAN-2020

DEFINITION	pSC101-P_araB_-sgRNA-Donor-T3

ACCESSION	pSC101-P_araB_-sgRNA-Donor-T3

KEYWORDS

SOURCE

ORGANISM

FEATURES	Location/Qualifiers

CDS	complement(1..951)

/label="rep101"

/note="=replication protein for the pSC101 origin"

rep_origin	999..1221

/label="pSC101 ori"

/note=" replication origin"

misc_feature	1843..1860

/label="I-SceI"

/note="recognition sequence of homing endonuclease I-SceI"

promoter	2006..2171

/label="araBAD promoter"

CDS	complement(2172..3032)

/label="AmpR"

/note="ampicillin resistance gene"

promoter	complement(3033..3137)

/label="AmpR promoter"

misc_RNA	3608..3683

/label="gRNA scaffold"

/note="guide RNA scaffold for the CRISPR/Cas9 system"

terminator	3744..3815

/label="rrnB T1 terminator"

ORIGIN

1 TCAGATCCTT CCGTATTTAG CCAGTATGTT CTCTAGTGTG GTTCGTTGTT TTTGCGTGAG

61 CCATGAGAAC GAACCATTGA GATCATACTT ACTTTGCATG TCACTCAAAA ATTTTGCCTC

121 AAAACTGGTG AGCTGAATTT TTGCAGTTAA AGCATCGTGT AGTGTTTTTC TTAGTCCGTT

181 ACGTAGGTAG GAATCTGATG TAATGGTTGT TGGTATTTTG TCACCATTCA TTTTTATCTG

241 GTTGTTCTCA AGTTCGGTTA CGAGATCCAT TTGTCTATCT AGTTCAACTT GGAAAATCAA

301 CGTATCAGTC GGGCGGCCTC GCTTATCAAC CACCAATTTC ATATTGCTGT AAGTGTTTAA

361 ATCTTTACTT ATTGGTTTCA AAACCCATTG GTTAAGCCTT TTAAACTCAT GGTAGTTATT

421 TTCAAGCATT AACATGAACT TAAATTCATC AAGGCTAATC TCTATATTTG CCTTGTGAGT

481 TTTCTTTTGT GTTAGTTCTT TTAATAACCA CTCATAAATC CTCATAGAGT ATTTGTTTTC

541 AAAAGACTTA ACATGTTCCA GATTATATTT TATGAATTTT TTTAACTGGA AAAGATAAGG

601 CAATATCTCT TCACTAAAAA CTAATTCTAA TTTTTCGCTT GAGAACTTGG CATAGTTTGT

661 CCACTGGAAA ATCTCAAAGC CTTTAACCAA AGGATTCCTG ATTTCCACAG TTCTCGTCAT

721 CAGCTCTCTG GTTGCTTTAG CTAATACACC ATAAGCATTT TCCCTACTGA TGTTCATCAT

781 CTGAGCGTAT TGGTTATAAG TGAACGATAC CGTCCGTTCT TTCCTTGTAG GGTTTTCAAT

841 CGTGGGGTTG AGTAGTGCCA CACAGCATAA AATTAGCTTG GTTTCATGCT CCGTTAAGTC

901 ATAGCGACTA ATCGCTAGTT CATTTGCTTT GAAAACAACT AATTCAGACA TACATCTCAA

961 TTGGTCTAGG TGATTTTAAT CACTATACCA ATTGAGATGG GCTAGTCAAT GATAATTACT

1021 AGTCCTTTTC CTTTGAGTTG TGGGTATCTG TAAATTCTGC TAGACCTTTG CTGGAAAACT

1081 TGTAAATTCT GCTAGACCCT CTGTAAATTC CGCTAGACCT TTGTGTGTTT TTTTTGTTTA

1141 TATTCAAGTG GTTATAATTT ATAGAATAAA GAAAGAATAA AAAAAGATAA AAAGAATAGA

1201 TCCCAGCCCT GTGTATAACT CACTACTTTA GTCAGTTCCG CAGTATTACA AAAGGATGTC

1261 GCAAACGCTG TTTGCTCCTC TACAAAACAG ACCTTAAAAC CCTAAAGGCT TAAGTAGCAC

1321 CCTCGCAAGC TCGGTTGCGG CCGCAATCGG GCAAATCGCT GAATATTCCT TTTGTCTCCG

1381 ACCATCAGGC ACCTGAGTCG CTGTCTTTTT CGTGACATTC AGTTCGCTGC GCTCACGGCT

1441 CTGGCAGTGA ATGGGGGTAA ATGGCACTAC AGGCGCCTTT TATGGATTCA TGCAAGGAAA

1501 CTACCCATAA TACAAGAAAA GCCCGTCACG GGCTTCTCAG GGCGTTTTAT GGCGGGTCTG

1561 CTATGTGGTG CTATCTGACT TTTTGCTGTT CAGCAGTTCC TGCCCTCTGA TTTTCCAGTC

1621 TGACCACTTC GGATTATCCC GTGACAGGTC ATTCAGACTG GCTAATGCAC CCAGTAAGGC

1681 AGCGGTATCA TCAACGGGGT CTGACGCTCA GTGGAACGAA AACTCACGTT AAGGGATTTT

1741 GGTCATGAGA TTATCAAAAA GGATCTTCAC CTAGATCCTT TTAAATTAAA AATGAAGTTT

1801 TAAATCAATC TAAAGTATAT ATGAGTAAAC TTGGTCTGAC AGTAGGGATA ACAGGGTAAT

1861 ACTTTTCATA CTCCCGCCAT TCAGAGAAGA AACCAATTGT CCATATTGCA TCAGACATTG

1921 CCGTCACTGC GTCTTTTACT GGCTCTTCTC GCTAACCAAA CCGGTAACCC CGCTTATTAA

1981 AAGCATTCTG TAACAAAGCG GGACCAAAGC CATGACAAAA ACGCGTAACA AAAGTGTCTA

2041 TAATCACGGC AGAAAAGTCC ACATTGATTA TTTGCACGGC GTCACACTTT GCTATGCCAT

2101 AGCATTTTTA TCCATAAGAT TAGCGGATCC TACCTGACGC TTTTTATCGC AACTCTCTAC

2161 TGTTTCTCCA TTTACCAATG CTTAATCAGT GAGGCACCTA TCTCAGCGAT CTGTCTATTT

2221 CGTTCATCCA TAGTTGCCTG ACTCCCCGTC GTGTAGATAA CTACGATACG GGAGGGCTTA

2281 CCATCTGGCC CCAGTGCTGC AATGATACCG CGAGACCCAC GCTCACCGGC TCCAGATTTA

2341 TCAGCAATAA ACCAGCCAGC CGGAAGGGCC GAGCGCAGAA GTGGTCCTGC AACTTTATCC

2401 GCCTCCATCC AGTCTATTAA TTGTTGCCGG GAAGCTAGAG TAAGTAGTTC GCCAGTTAAT

2461 AGTTTGCGCA ACGTTGTTGC CATTGCTACA GGCATCGTGG TGTCACGCTC GTCGTTTGGT

2521 ATGGCTTCAT TCAGCTCCGG TTCCCAACGA TCAAGGCGAG TTACATGATC CCCCATGTTG

2581 TGCAAAAAAG CGGTTAGCTC CTTCGGTCCT CCGATCGTTG TCAGAAGTAA GTTGGCCGCA

2641 GTGTTATCAC TCATGGTTAT GGCAGCACTG CATAATTCTC TTACTGTCAT GCCATCCGTA

2701 AGATGCTTTT CTGTGACTGG TGAGTACTCA ACCAAGTCAT TCTGAGAATA GTGTATGCGG

2761 CGACCGAGTT GCTCTTGCCC GGCGTCAATA CGGGATAATA CCGCGCCACA TAGCAGAACT

2821 TTAAAAGTGC TCATCATTGG AAAACGTTCT TCGGGGCGAA AACTCTCAAG GATCTTACCG

2881 CTGTTGAGAT CCAGTTCGAT GTAACCCACT CGTGCACCCA ACTGATCTTC AGCATCTTTT

2941 ACTTTCACCA GCGTTTCTGG GTGAGCAAAA ACAGGAAGGC AAAATGCCGC AAAAAAGGGA

3001 ATAAGGGCGA CACGGAAATG TTGAATACTC ATACTCTTCC TTTTTCAATA TTATTGAAGC

3061 ATTTATCAGG GTTATTGTCT CATGAGCGGA TACATATTTG AATGTATTTA GAAAAATAAA

3121 CAAATAGGGG TTCCGCGGCA CAGATGCGTA AGGAGAAAAT ACCGCATCAG GCGCCATTCG

3181 CCATTCAGGC TGCGCAACTG TTGGGAAGGG CGATCGGTGC GGGCCTCTTC GCTATTACGC

3241 CAGCTGGCGA AAGGGGGATG TGCTGCAAGG CGATTAAGTT GGGTAACGCC AGGGTTTTCC

3301 CAGTCACGAC GTTGTAAAAC GACGGCCAGT GCCAAGCTTG CATGCCTGCA GGTCGACTCT

3361 AGAGGATCCC CGGGTACCGA GCTCGAATTC GTAATCATGT CATAGCTGTT TCCTGTGTGA

3421 AATTGTTATC CGCTCACAAT TCCACACAAC ATACGAGCCG GAAGCATAAA GTGTAAAGCC

3481 TGGGGTGCCT AATGAGTGAG CTAACTCACA TTAATTGCGT TGCGCTCACT GCCCGCTTTC

3541 CAGTCGGGAA ACCTGTCATA ACACCGTGCG TGTTGACTAT TTTACCTCTG GCGGTGATAA

3601 TGGTTGCGTT TTAGAGCTAG AAATAGCAAG TTAAAATAAG GCTAGTCCGT TATCAACTTG

3661 AAAAAGTGGC ACCGAGTCGG TGCTTAGCAT CCAAACTCGA GTAAGGATCA TTAAGGATCC

3721 CATGGTACGC GTGCTAGAGG CATCAAATAA AACGAAAGGC TCAGTCGAAA GACTGGGCCT

3781 TTCGTTTTAT CTGTTGTTTG TCGGTGAACG CTCTCCTGAG TAGGACAAAT CCGCCCCATG

3841 GGTATGGACA GTTTTCCCTT TGATATGTAA CGGTGAACAG TTGTTCTACT TTTGTTTGTT

3901 AGTCTTGATG CTTCACTGAT AGATACAAGA GCCATAAGAA CC

//

**References**

1.	B. P. Anton, E. A. Raleigh. Complete genome sequence of NEB 5-alpha, a derivative of *Escherichia coli* K-12 DH5α*.* Genome Announc., 2016, 4: e01245-16

2.	K. Hayashi, N. Morooka, Y. Yamamoto, K. Fujita, K. Isono, S. Choi, et al. Highly accurate genome sequences of *Escherichia coli* K-12 strains MG1655 and W3110*.* Mol. Syst. Biol., 2006, 2: 2006.0007

3.	J. Norrander, T. Kempe, J. Messing. Construction of improved M13 vectors using oligodeoxynucleotide-directed mutagenesis*.* Gene, 1983, 26: 101–106
